# High-pressure Golgi neuronal staining for X-ray whole-brain imaging

**DOI:** 10.1101/2025.01.04.631279

**Authors:** Qiaowei Tang, Xiaoqing Cai, Yu Liu, Yu Zhang, Xin Yan, Feng Zhou, Jichao Zhang, Qian Li, Ke Li, Biao Deng, Lihua Wang, Jiang Li, Ying Zhu, Chunhai Fan, Jun Hu

**Author notes:** These authors contributed equally: Qiaowei Tang, Xiaoqing Cai, Yu Liu, Yu Zhang.

## Abstract

Whole-brain imaging has revolutionized neuroscience research, providing comprehensive insights into neural networks across the entire brain. This powerful approach has greatly advanced our understanding of brain functions and the mechanisms underlying various diseases. One primary challenge in whole-brain imaging technology is to achieve high-resolution observation of neural networks at large scales. Although Golgi method allows labeling random neurons in their entirety in the brain, visualizing individual dendritic trees, and tracing long-distance axonal projections, the lengthy processing time pose a limit on its use, i.e., staining a mouse whole-brain sample of just 300 mm³ takes over two weeks. Here, we developed a rapid staining technique for whole-brain neurons using high-pressure assisted Golgi (HP Golgi). This method significantly reduced the staining time for mouse whole-brain neurons from 16 days to only 4 days. We demonstrated the broad applicability of the HP Golgi method across various model organisms, achieving whole-brain neuronal staining in zebrafish, mice, and rats. Further, we successfully performed rapid staining of hippocampal neurons in an intact pig brain, which is difficult to achieve with the classic Golgi-Cox method. We also demonstrated that the combination of the HP Golgi method with synchrotron-based X-ray microscopy for high-resolution imaging of whole-brain neurons in mice. This HP Golgi method enables fast and high-resolution neuronal imaging in large model organisms, showcasing its broad applicability for diverse applications.

## Introduction

Whole-brain imaging techniques are indispensable for unraveling the intricate neural networks underlying brain function and developing effective therapeutic strategies for neurological disorders ^1^. For instance, electron microscopy (EM) has been utilized to reconstruct the whole-brain connectome of *Drosophila melanogaster* at synaptic resolution ^2^, while fluorescence micro-optical sectioning tomography (fMOST) has facilitated single-neuron-level mapping of hippocampal projectomes in mice ^3^. Luo et al. expanded the utility of fMOST by constructing a comprehensive whole-brain atlas of the cholinergic system at the single-neuron level, providing a crucial resource for exploring neurological conditions such as dementia, sleep disorders, and cognitive deficits associated with cholinergic dysfunction ^4^. Whole-brain imaging integrates diverse neuronal staining and labeling techniques with state-of-the-art imaging technologies. Among these, mesoscopic imaging approaches currently offer the most practical solutions for single-neuron studies at the whole-brain scale. Key methodologies include fMOST, volumetric imaging with synchronized on-the-fly scanning and readout (VISoR), and serial two-photon tomography (STPT) ^5–8^.

Among these imaging approaches, achieving effective staining or labeling of large brain tissues remains a critical challenge. One strategy employs viral tools combined with tissue sectioning, allowing for serial staining of individual sections or simultaneous staining during sectioning. By integrating advanced section registration techniques, this approach enables high-resolution three-dimensional reconstruction of entire brains. Using these methods, researchers have successfully performed whole-brain imaging at subcellular resolution in mice and mapped neuronal projectomes, such as prefrontal cortex neurons, paraventricular hypothalamic (PVH) oxytocin neurons, and axonal fiber projections in rhesus monkeys ^9–13^. However, structural damage at section boundaries often compromises neuronal integrity and overall brain architecture, introducing deformation and reducing the fidelity of experimental results ^14^. Additionally, viral labeling techniques typically require 2–4 weeks for sufficient expression, further limiting their efficiency ^15, 16^. Another promising strategy combines tissue-clearing techniques with whole-brain immunostaining. These methods mitigate the issues associated with extensive sectioning; however, achieving uniform staining in large brain samples is time-consuming and labor-intensive. For instance, clearing and staining a 300 mm³ mouse brain using techniques such as iDISCO+ or CUBIC-HistoVision requires 18 days and 2 weeks, respectively ^17–19^. These limitations pose significant challenges to the high-throughput application of whole-brain imaging.

In 1873, the Italian scientist Camillo Golgi developed the Golgi method, a groundbreaking technique that used chromate-silver nitrate staining to visualize the fundamental structures of neurons— including axons, dendrites, and dendritic spines—under an optical microscope ^20, 21^. This method was later refined by the Spanish scientist Santiago Ramón y Cajal, who enhanced its staining stability ^22^. To this day, the Golgi staining method remains a critical tool for studying neuronal morphology ^23^. In 2010, the team led by Qingming Luo applied the Golgi-Cox method for whole-brain staining in mice and combined it with their self-developed Micro-Optical Sectioning Tomography (MOST) imaging system to generate three-dimensional imaging data of the entire mouse brain ^5^. However, this technique revealed a key limitation. While the imaging phase was completed in 10 days, neuronal staining took 180 days, creating an 18-fold disparity and becoming a major bottleneck in whole-brain imaging data acquisition. In 2014, Rosoklija and colleagues introduced modifications to the Golgi-Cox method, such as gentle agitation on a slowly rocking platform and reduced exposure to ammonia, enabling the staining depth of 1 cm in human brain tissue sections within 6 weeks. These improvements yielded high-quality morphological images of cortical neurons under optical microscopy ^24^. More recently, several commercial kits based on the Golgi-Cox method, including the Hito Golgi-Cox OptimStain™ Kit and FD Rapid GolgiStain™ Kit, have been developed to further reduce staining time ^25, 26^. Despite these advancements, staining an entire mouse brain using these kits still requires 2–3 weeks, underscoring the need for faster and more efficient staining techniques.

Inspired by the application of high-pressure techniques in histology to reduce tissue processing time and expedite pathological diagnoses ^27, 28^, we developed a novel high-pressure-assisted Golgi staining method for rapid whole-brain neuronal visualization. We systematically investigated the effects of variables such as temperature, pressure, and duration on staining efficiency. Our findings reveal that applying a pressure of 9 MPa reduces the staining time for mouse whole-brain neurons to just one-quarter of that required by commercial kits. We further validated the practicality of this rapid staining method across multiple model organisms and functional brain regions. By integrating this method with synchrotron-based X-ray microscopy, we achieved rapid neuronal staining and high-resolution imaging of the entire mouse brain (Fig. 1).

**Fig. 1.**
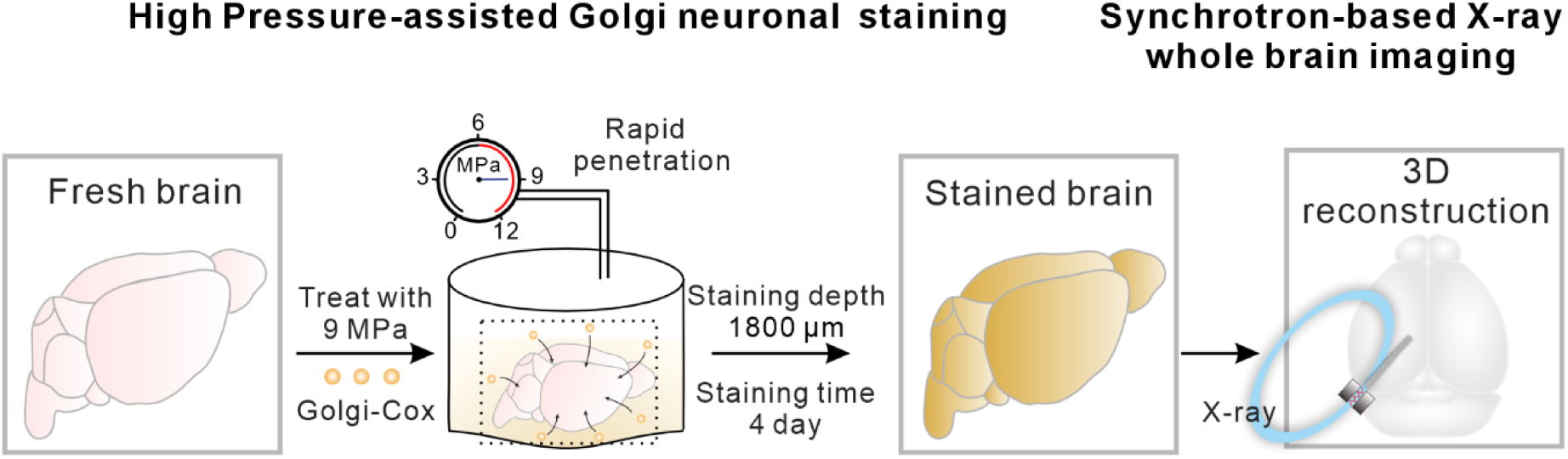
Schematic illustrating high-pressure Golgi neuronal staining for synchrotron-based X-ray whole-brain imaging. A fresh mouse brain was stained using the HP Golgi method for 4 days (37°C, 9 MPa), yielding staining results comparable to those of the classical Golgi-Cox method, which requires 16 days. This technique enabled neuronal staining to a depth of 1800 μm within 2 hours. After staining, the entire brain was imaged using a synchrotron-based X-ray microscope, facilitating high-resolution whole-brain imaging.

## Results

### Establishment of high-pressure Golgi neuronal staining method

An ideal method for labeling whole-brain neurons in large-scale model organisms should combine high contrast, rapid processing, and precise, sensitive detection compatible with microscopy techniques. When designing our approach, we addressed the well-known limitations of the Golgi-Cox method, particularly its slow penetration of staining solutions into brain tissue. For example, staining the relatively small mouse brain (∼300 mm³) often requires several weeks ^29^. To overcome these constraints, we developed a high-pressure-assisted Golgi (HP-Golgi) method to accelerate the staining process. In our approach, whole mouse brains were immersed in a Golgi staining solution containing mercuric chloride, potassium dichromate, and potassium chromate and subjected to high-pressure device (Fig. 2a). A major limitation of the classical Golgi-Cox method is its incompatibility with fixatives such as paraformaldehyde, which leads to incomplete staining of deeper brain regions over short timeframes, increasing the risk of tissue autolysis ^30^. The HP-Golgi method enhances the penetration of the staining solution into brain tissue under high pressure, ensuring rapid interaction of the solution with internal brain structures. This not only significantly reduces the staining duration but also maximizes the preservation of tissue integrity.

**Fig. 2.**
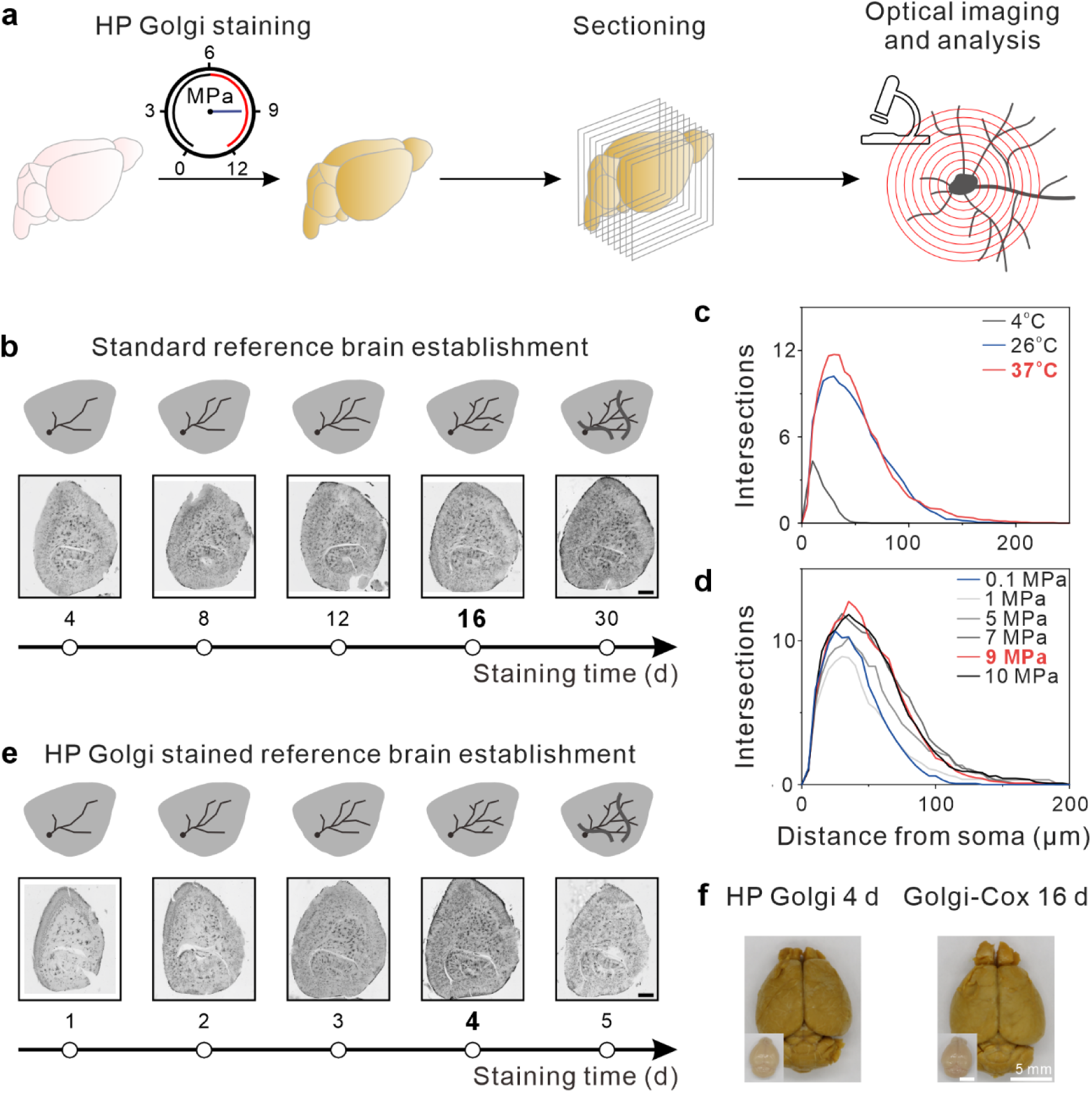
Establishment of the high-pressure Golgi neuronal staining method. **a,** Workflow of the HP Golgi method. Fresh mouse brains were stained under 9 MPa, followed by sectioning and optical microscopy analysis. **b,** Standard reference brain establishment. **Top:** Schematic of neurons and blood vessels in a brain slice; **Bottom:** Panoramic images of the brain slice. Whole mouse brains were stained using the Golgi-Cox method for 4, 8, 12, 16, and 30 days. As staining duration increased, neuronal structures became progressively more complete. Notably, significant vascular staining was observed in the panoramic image at 30 days. Scale bar: 1 mm. **c,** Sholl analysis of cortical neurons stained at different temperatures. At atmospheric pressure, whole mouse brains were stained at 4L, 26L, and 37L for 4 days. Following sectioning, Sholl analysis was performed on SS neurons. **d,** Sholl analysis of cortical neurons stained at different pressures. At 37L, whole mouse brains were stained under 0.1 MPa, 1 MPa, 5 MPa, 7 MPa, 9 MPa, and 10 MPa for 1 day. After sectioning, Sholl analysis was performed on SS neurons. **e,** HP Golgi standard reference brain establishment. **Top:** Schematic of neurons and blood vessels in a brain slice; **Bottom:** Panoramic images of the brain slice. Whole mouse brains were stained using the HP Golgi method for 1, 2, 3, 4, and 5 days. As staining duration increased, neuronal structures became progressively more complete. Significant vascular staining was observed at 5 days. Scale bar: 1 mm. **f,** Comparison of brain images before and after staining. **Left:** HP Golgi method for 4 days; **Right:** Golgi-Cox method for 16 days.

To evaluate whether the high-pressure-assisted Golgi (HP-Golgi) method enhances neuronal staining efficiency, we used the classical Golgi-Cox method as a reference. In a standard staining experiment, whole mouse brains were dissected, immersed in Golgi staining solution, and processed in the dark for varying durations. Visual inspection revealed a gradual color change from bright yellow to brownish yellow as staining time increased (Supplementary Fig. 1). Sagittal sections of the stained brain tissue were prepared using a cryostat and imaged under an optical microscope. For staining durations of 1–16 days, clear neuronal morphology was observed, accompanied by minimal blood vessel staining, resulting in a high neuron-to-blood vessel staining ratio (Fig. 2b and Supplementary Fig. 2). Sholl analysis of neurons in the somatosensory cortex (SS) indicated that with increasing staining time, neurons exhibited more proximal intersections (20–60 μm) and extended dendritic branches at distal locations (>150 μm) (Supplementary Fig. 3). These findings suggest progressive improvement in neuronal structure visibility and completeness over time, reaching optimal results within 16 days. For staining durations exceeding 16 days, observations of brain sections revealed a marked increase in blood vessel staining, particularly in neocortical regions such as the visual cortex (VIS), somatosensory cortex (SS), and somatomotor cortex (MO) (Fig. 2b and Supplementary Fig. 2). This excessive vascular staining complicated the visualization and segmentation of neuronal structures. Based on these results, we identified 16 days as the optimal staining duration for the Golgi-Cox method in whole mouse brains. This balance minimizes vascular staining while achieving comprehensive neuronal labeling, enabling effective structural analysis of neuronal networks.

Next, we investigated the influence of temperature on the Golgi staining process by conducting experiments at 4°C, 26°C, 37°C, and 60°C under ambient pressure and light-protected conditions for 4 days. Visual inspection revealed that as the staining temperature increased, the color of the mouse brain shifted progressively from bright yellow to dark brown. Notably, brains stained at 60°C displayed a deeper color than those processed for 30 days under standard Golgi-Cox staining conditions (Supplementary Fig. 4). To evaluate neuronal morphology, sagittal sections of the stained brains were prepared and imaged using an optical microscope. At 4°C, 26°C, and 37°C, neuronal structures were clearly discernible, with minimal vascular staining (Supplementary Fig. 4). However, at 4°C, dendritic branches were shorter compared to those observed at 26°C and 37°C, suggesting reduced staining completeness at lower temperatures. In contrast, at 60°C, neuronal structures were poorly resolved, and vascular staining was significantly elevated, indicating a reduced neuron-to-blood vessel staining ratio. Panoramic images revealed tissue damage in samples stained at 60°C, highlighting the adverse effects of excessive temperatures on tissue integrity (Supplementary Fig. 4). To quantify these findings, we performed Sholl analysis of neurons in the SS under different temperature conditions. Neurons stained at 37°C and 26°C exhibited greater complexity, as indicated by a higher number of intersections, compared to those at 4°C (Fig. 2c). Specifically, neurons in the 37°C group demonstrated approximately 11% more proximal intersections (40 μm from the soma) and a 27% increase in maximum dendritic length compared to those in the 26°C group. These results identify 37°C as the optimal temperature for achieving high neuronal staining specificity and morphological complexity while maintaining tissue integrity.

At the optimal temperature of 37°C, we further assessed the effect of high pressure on neuronal staining in mouse brains. The staining experiments were conducted under pressures of 0.1 MPa (atmospheric pressure), 1 MPa, 5 MPa, 7 MPa, 9 MPa, and 10 MPa, followed by tissue sectioning and imaging under an optical microscope (Supplementary Fig. 5). Panoramic images revealed no discernible tissue damage or significant background staining across the pressure conditions, indicating the preservation of tissue integrity (Supplementary Fig. 5). Magnified views of the somatosensory cortex (SS) showed clear neuronal morphology with minimal vascular staining across all pressure groups (Supplementary Fig. 5). Sholl analysis of neuronal complexity revealed pressure-dependent effects. At proximal regions (e.g., 40 μm), neurons in the 1 MPa group exhibited a 12% decrease in mean intersection counts compared to the 0.1 MPa group, while the maximum dendritic branch length increased by approximately 19%. For the 5 MPa, 7 MPa, and 9 MPa groups, the mean intersection counts increased by 2%, 17%, and 31%, respectively, with corresponding increases in maximum dendritic branch length of 50%, 42%, and 50%. These results indicate that neuronal complexity was significantly enhanced at 9 MPa relative to atmospheric pressure. However, at 10 MPa, the intersection count increased by 21% compared to atmospheric pressure, but the maximum dendritic branch length plateaued at 195 μm, showing no further improvement (Fig. 2d). These findings demonstrate that a pressure of 9 MPa significantly enhances the Golgi-Cox staining of mouse brain neurons, achieving improved neuronal complexity and dendritic visualization without compromising tissue integrity. Thus, 9 MPa was selected as the optimal pressure for subsequent experiments.

Using the optimized conditions of 37°C and 9 MPa, we investigated the effect of staining duration on the HP-Golgi method. Staining was performed for 1, 2, 3, 4, and 5 days, and brain sections were analyzed via optical microscopy (Fig. 2e). Panoramic images revealed that neuronal morphology was clearly visible with minimal vascular staining for staining durations of 1–4 days, resulting in a high neuron-to-blood vessel staining ratio (Fig. 2e). However, at 5 days, vascular staining increased markedly, particularly in regions such as the motor cortex, complicating neuronal visualization and segmentation due to reduced neuron-to-vessel specificity (Fig. 2e). Sholl analysis of neurons in the SS after 1–4 days of staining demonstrated progressive improvements in neuronal complexity. With increasing staining time, neurons exhibited a greater number of intersections at proximal distances (20–60 μm) and more extended dendritic branches at distal locations (>150 μm). At 40 μm, the mean intersection counts were 7.7, 10.0, 12.0, and 13.5 for 1, 2, 3, and 4 days of staining, respectively (Supplementary Fig. 6). These results indicate that a 4-day staining duration achieves the most complete neuronal staining under high-pressure conditions while maintaining low background interference. In summary, a staining duration of 4 days under 37°C and 9 MPa provides optimal conditions for achieving high-quality neuronal staining with clear morphology and minimal background, facilitating subsequent imaging and analysis.

We compared neuronal staining outcomes achieved with the HP-Golgi method (4 days) to those obtained using the classical Golgi-Cox method (16 days). Both methods yielded whole mouse brains with comparable coloration (Fig. 2f). Sagittal sections at a depth of 2000 μm were imaged under an optical microscope, capturing panoramic views and magnified regions of interest, including the somatosensory cortex (SS) and hippocampus (Fig. 3a, b). The imaging results demonstrated that neuronal staining across the whole brain was similar between the two approaches. Notably, the HP-Golgi method resulted in significantly reduced vascular staining during the shorter 4-day staining period, particularly in the SS region (Fig. 3a, b). Sholl analysis of SS neurons further confirmed that neuronal complexity metrics, such as dendritic intersections, were comparable between the two methods (Supplementary Fig. 7). This indicates equivalent completeness of neuronal staining in the SS region. In summary, the HP-Golgi method produces neuronal staining results equivalent to those of the Golgi-Cox method while reducing the staining duration from 16 days to just 4 days. This fourfold reduction in time underscores the HP-Golgi method as a rapid and efficient alternative for whole-brain neuronal staining

**Fig. 3.**
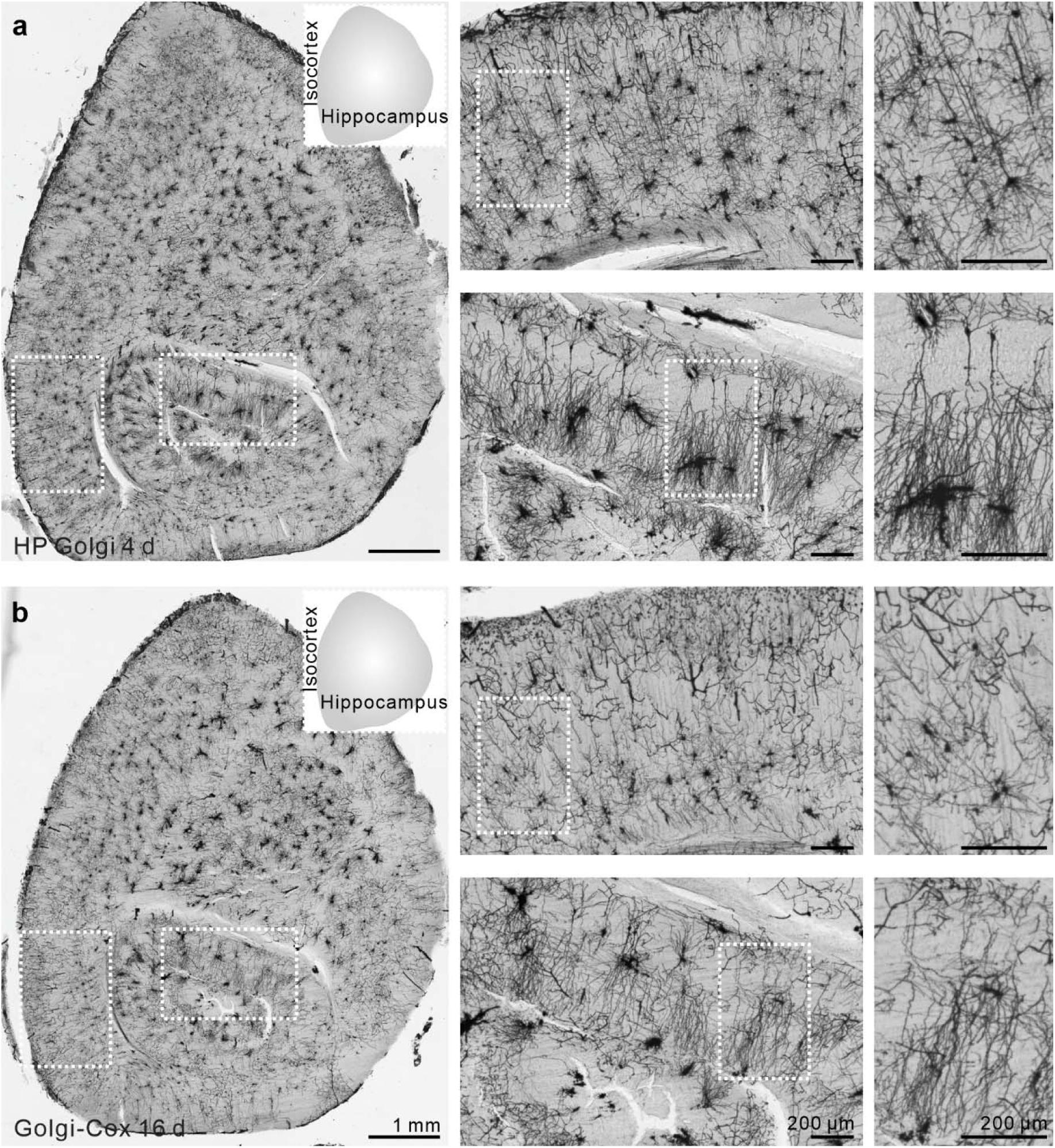
Comparison of panoramic images of brain neurons in C57BL/6 mice using optical microscopy with HP Golgi and classic Golgi-Cox staining methods. **a,** Panoramic image and localized magnification of a mouse brain section stained using the HP Golgi method for 4 days. **Left:** Panoramic image with schematic highlighting the hippocampus and isocortex. Scale bar: 1 mm. **Right:** Magnification of specific brain regions. **Top:** Isocortex magnification. Scale bar: 200 μm. **Bottom:** Hippocampus magnification. Scale bar: 200 μm. **b,** Panoramic image and localized magnification of a mouse brain section stained using the Golgi-Cox method for 16 days. **Left:** Panoramic image with schematic highlighting the hippocampus and isocortex. Scale bar: 1 mm. **Right:** Magnification of specific brain regions. **Top:** Isocortex magnification. Scale bar: 200 μm. **Bottom:** Hippocampus magnification. Scale bar: 200 μm.

### Mechanism analysis of high-pressure-empowered fast Golgi neuronal staining

To investigate the reasons behind the reduced staining time with the HP Golgi method in whole mouse brains, we assessed the penetration behavior of the Golgi staining solution within mouse brain tissue. Whole mouse brains were stained using both the HP Golgi and Golgi-Cox methods for 2 hours, followed by continuous sagittal sectioning at 200 μm intervals. Panoramic images were acquired for each section to compare the penetration characteristics of the two methods under identical staining conditions. Figures 4a and 4b illustrate the imaging results for sequential sections stained with the HP Golgi and Golgi-Cox methods, respectively. The HP Golgi method demonstrated effective neuronal staining in the central regions of sections up to a depth of 1800 μm (Fig. 4a). At 2000 μm, neuronal staining was no longer detectable in the central areas, except in the hippocampus, where staining persisted. This effect likely arises from tissue gaps that allow deeper penetration of the staining solution into the hippocampus, suggesting that the effective staining depth for the HP Golgi method is approximately 1800 μm. In contrast, the Golgi-Cox method revealed neuronal staining in the central regions only up to 600 μm, with staining becoming undetectable at depths beyond 800 μm (Fig. 4b). These results demonstrate that the HP Golgi method enables faster penetration of the staining solution, allowing for deeper brain regions to be stained in a shorter period compared to the classical Golgi-Cox method.

**Fig. 4.**
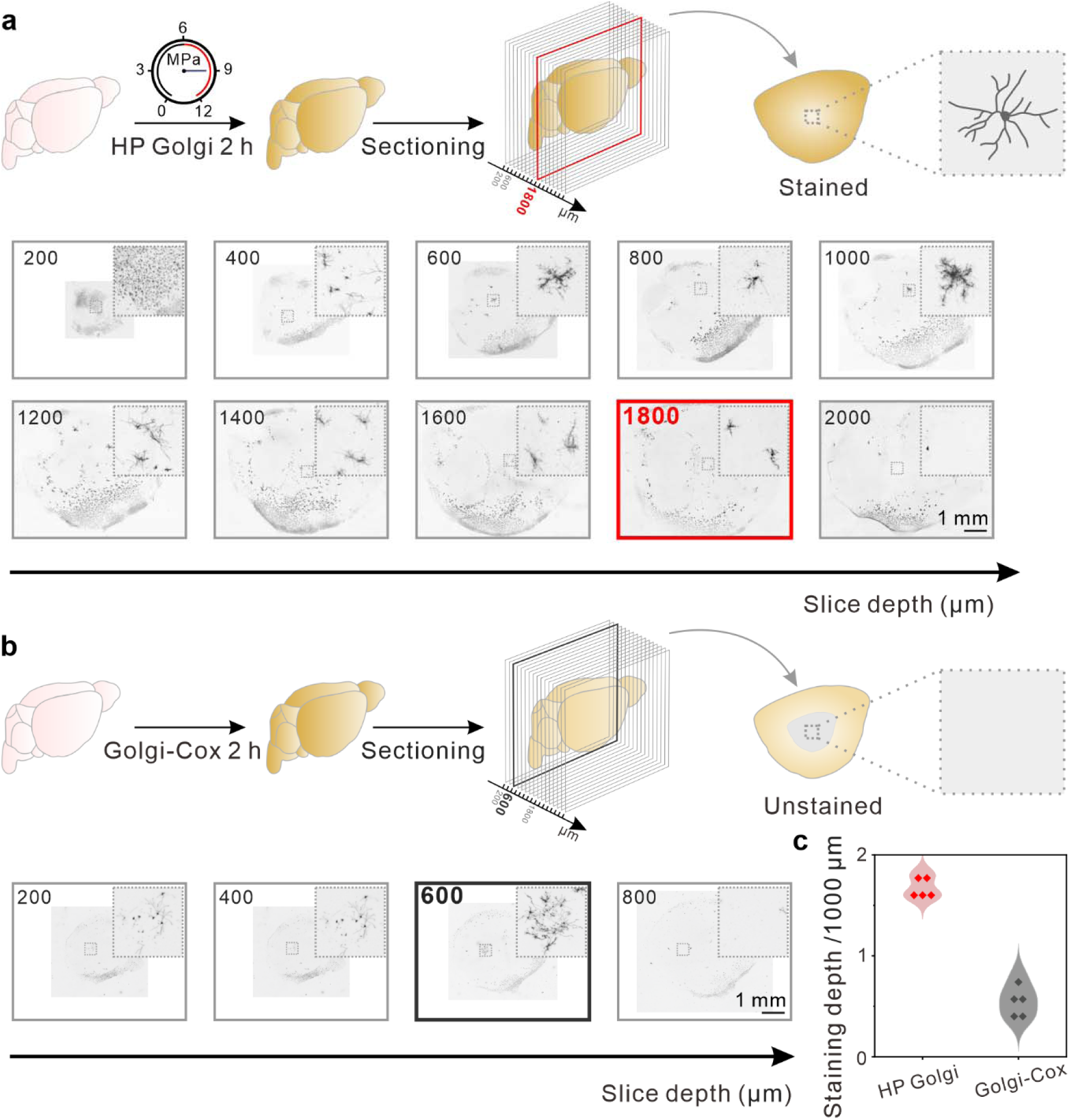
Mechanism analysis of high-pressure fast Golgi neuronal staining. **a,** Scheme and serial section images of slices stained using the HP Golgi method for 2 hours. **Top:** Schematic diagram showing the staining depth. The HP Golgi method achieved a staining depth of 1800 μm within 2 hours. **Bottom:** Serial section images. Neuronal staining was visible in the central regions of the sections at depths ranging from 200 to 1800 μm, but no staining was observed at a depth of 2000 μm in the central regions of the brain tissue. Scale bar: 1 mm. **b,** Scheme and serial section images of slices stained using the Golgi-Cox method for 2 hours. **Top:** Schematic diagram showing the staining depth. The Golgi-Cox method achieved a staining depth of 600 μm within 2 hours. **Bottom:** Serial section images. Neuronal staining was visible in the central regions of the sections at depths ranging from 200 to 600 μm, but no staining was observed at a depth of 800 μm in the central regions of the brain tissue. Scale bar: 1 mm. **c,** Violin plot of the statistical analysis of staining depth. For the HP Golgi method, the average staining depth was 1680 μm, while for the Golgi-Cox method, the average staining depth was 560 μm.

Next, we performed a statistical analysis of staining depths in two groups of mouse brains (five mice per group). After 2 hours of staining, the HP Golgi method achieved an average staining depth of 1680 μm, compared to 560 μm with the Golgi-Cox method (Fig. 4c), representing a threefold enhancement. This improvement underscores the efficacy of the HP Golgi method in facilitating rapid penetration of staining solutions into brain tissue. Traditional fixatives such as formaldehyde are known to increase glial cell staining, which contributes to elevated background signals in tissue samples. To mitigate this issue, neither the HP Golgi nor the classical Golgi-Cox method employed fixatives. While this approach effectively reduces background staining, it introduces the risk of tissue autolysis, potentially compromising neuronal morphology. Notably, the Golgi staining solution contains potassium dichromate, which imparts mild fixation properties while minimizing the background staining of tissue sections ^31^. The HP Golgi method further enhances staining efficiency by promoting rapid penetration of the solution into brain tissue. This accelerated infiltration preserves the internal morphological structure and reduces the likelihood of autolysis, thereby improving the efficiency and reliability of whole-brain neuronal staining.

### Broad applicability of HP Golgi neuronal staining method

To assess the versatility of our method across animal models with diverse neuronal morphologies and brain structures, we conducted neuronal staining in representative models, including zebrafish, rats, and pigs ^11, 32–34^. Initially, we applied the classic Golgi-Cox method to zebrafish brains, observing samples at 1, 2, 3, 4, 5, and 6 days (Fig. 5a, Supplementary Fig. 8). Optical microscopy revealed clear neuronal morphology at all time points; however, by day 6, increased vascular staining hindered analysis. Sholl analysis indicated that extended staining durations resulted in more complete neuronal morphologies (Supplementary Fig. 9). Subsequently, we applied the HP Golgi method for 1, 2, and 3 days of staining, which also yielded distinct neuronal structures (Fig. 5b, Supplementary Fig. 10). Sholl analysis showed that 3 days of HP Golgi staining resulted in more complete neuronal structures compared to 1 or 2 days (Supplementary Fig. 11). Notably, 3-day HP Golgi staining outperformed the 5-day Golgi-Cox method, with approximately a 18% increase in the number of intersection points at the proximal site (20 μm from the soma) and approximately a 43% improvement in the length of the longest dendritic branches at the distal site (Fig. 5c). These findings demonstrate that the HP Golgi method significantly reduces the staining time for zebrafish brains, requiring only 60% of the time compared to the classic Golgi-Cox method.

**Fig. 5.**
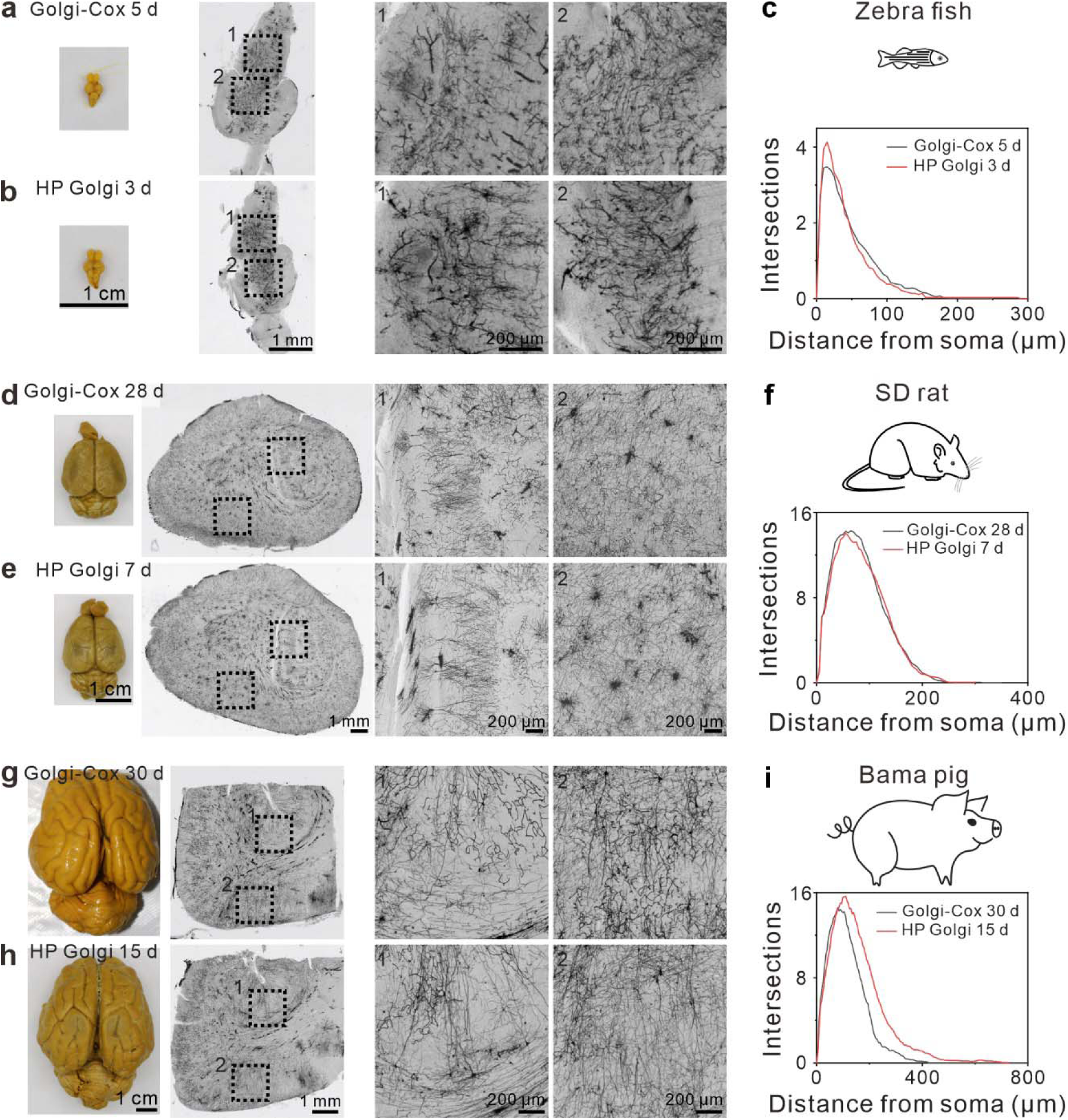
Broad applicability of HP Golgi neuronal staining method. **a,** Zebrafish brain stained for 5 days using the Golgi-Cox method. **Left:** Image of the zebrafish brain after staining. Scale bar: 1 mm. **Right:** Panoramic image and localized magnification of the zebrafish brain section. Scale bar: 1 mm (panoramic image) and 200 μm (magnification). **b,** Zebrafish brain stained for 3 days using the HP Golgi method. **Left:** Image of the zebrafish brain after staining. Scale bar: 1 mm. **Right:** Panoramic image and localized magnification of the zebrafish brain section. Scale bar: 1 mm (panoramic image) and 200 μm (magnification). **c,** Sholl analysis of zebrafish brain neurons for different staining methods. **d,** Rat brain stained for 28 days using the Golgi-Cox method. **Left:** Image of the rat brain after staining. Scale bar: 1 mm. **Right:** Panoramic image and localized magnification of the rat brain section. Scale bar: 1 mm (panoramic image) and 200 μm (magnification). **e,** Rat brain stained for 7 days using the HP Golgi method. **Left:** Image of the rat brain after staining. Scale bar: 1 mm. **Right:** Panoramic image and localized magnification of the rat brain section. Scale bar: 1 mm (panoramic image) and 200 μm (magnification). **f,** Sholl analysis of rat brain neurons for different staining methods. **g,** Pig brain stained for 30 days using the Golgi-Cox method. **Left:** Image of the pig brain after staining. Scale bar: 1 mm. **Right:** Panoramic image and localized magnification of the pig brain section. Scale bar: 1 mm (panoramic image) and 200 μm (magnification). **h,** Pig brain stained for 15 days using the HP Golgi method. **Left:** Image of the pig brain after staining. Scale bar: 1 mm. **Right:** Panoramic image and localized magnification of the pig brain section. Scale bar: 1 mm (panoramic image) and 200 μm (magnification). **i,** Sholl analysis of pig brain neurons for different staining methods.

In the following experiment, we utilized the Golgi-Cox staining method to stain rat brains for various durations (4, 7, 14, 21, 28, and 35 days). Optical microscopy revealed clear neuronal morphology for staining periods ranging from 4 to 28 days, with some vascular staining observed (Fig. 5d, Supplementary Fig. 12). However, staining for 35 days resulted in a significant increase in blood vessel staining, suggesting that prolonged staining reduces the neuron-to-blood vessel staining ratio, which may interfere with the accurate observation of neuronal structures (Fig. 5d). This finding aligns with recommendations in the literature that Golgi-Cox staining should not exceed one month ^35^. Next, we applied the HP Golgi method to stain rat brains for 4, 7, and 14 days, and compared the outcomes with those from the Golgi-Cox group (Fig. 5e, Supplementary Fig. 13). Neuronal staining was evident at all three time points, with an increasing number of stained blood vessels observed at the 14-day mark. Sholl analysis of neurons in the sensorimotor (SS) region demonstrated that, with longer staining durations, the neuronal morphology became progressively more complete (Supplementary Figs. 14 and 15). Notably, neurons stained for 7 days using the HP Golgi method exhibited complexity comparable to that of neurons stained for 28 days with the Golgi-Cox method, with no significant differences observed between the two groups (Fig. 5f). These findings indicate that the HP Golgi method reduces the staining time for rat brain neurons to one-fourth of the duration required by the classical Golgi-Cox method.

We also performed neuronal staining on intact porcine brains. Given that the neuron-to-blood vessel staining ratio significantly decreases after 30 days, we employed the Golgi-Cox method for 30-day staining (Fig. 5g). Macroscopic examination revealed a uniformly yellow-stained brain. However, optical microscopy of brain sections only revealed discernible neuronal structures in the cortical layers, with no visible neurons in deeper regions. Subsequently, we applied the HP Golgi method for staining durations of 10, 15, and 20 days (Fig. 5h, Supplementary Fig. 16). At 20 days, vascular staining was notably increased, while at 15 days, neuronal structures in the hippocampal region—previously inaccessible with the classical Golgi-Cox method—were successfully visualized. Additionally, blood vessel staining was less pronounced at 15 days compared to the 30-day Golgi-Cox staining, suggesting that the HP Golgi method not only enhances neuronal staining contrast but also proves more effective for staining deeper brain regions. Sholl analysis further demonstrated that neurons stained for 15 days with the HP Golgi method exhibited greater morphological complexity than those stained for 30 days with the Golgi-Cox method or for 10 days with the HP Golgi method (Fig. 5i). Specifically, at the proximal site (100 μm), the average number of intersection points increased by approximately 10%, while the length of dendritic branches in the distal site increased by approximately 70%. These findings demonstrate the superior efficacy of the HP Golgi method for staining porcine hippocampal neurons in intact porcine brains, a task that is challenging with the classical Golgi-Cox method.

### Synchrotron-based X-ray visualization of whole-brain neurons using HP-Golgi staining

Having established the broad applicability of the HP Golgi method for neuronal staining across multiple model organisms, we employed synchrotron-based X-ray micro-computed tomography (Micro-CT) to acquire high-resolution imaging data of neurons from zebrafish, rats, and pigs at the 13 HB beamline of the Shanghai Synchrotron Radiation Facility (SSRF) (Fig. 6a). Imaging was performed with a voxel size of 0.65 μm and an X-ray energy of 14 keV. Three-dimensional reconstructions and high-magnification images of cortical neurons revealed typical neuronal morphology, including cell bodies and dendrites, across all species (Fig. 6b–6d and Supplementary Videos 1–3). These results validate the applicability of HP Golgi staining for Micro-CT imaging of brain neurons in diverse model organisms.

**Fig. 6.**
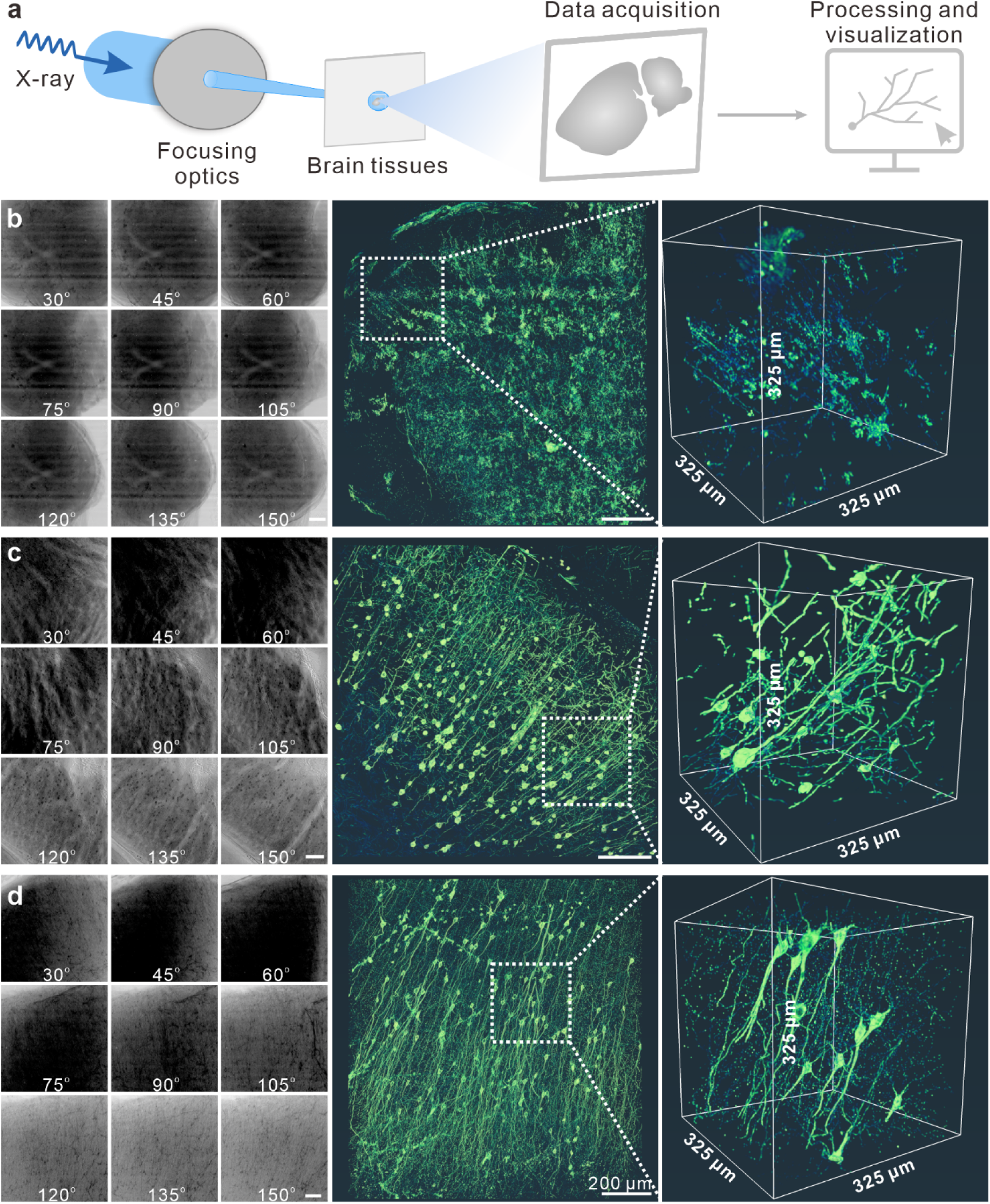
Synchrotron-based X-ray visualization of brain neurons in different animal models using the HP Golgi method. **a,** Schematic of synchrotron-based X-ray imaging and data processing. Brain samples were subjected to projection image acquisition using synchrotron-based X-ray imaging, followed by slice reconstruction of the projected images and three-dimensional visualization. **b,** Projections and three-dimensional visualization of the zebrafish brain. **Left:** Projections from different angles. Scale bar: 200 μm. **Center:** Three-dimensional visualization of the zebrafish brain. Scale bar: 200 μm. **Right:** Localized magnification of the center. **c,** Projections and three-dimensional visualization of the rat brain. **Left:** Projections from different angles. Scale bar: 200 μm. **Center:** Three-dimensional visualization of the rat brain. Scale bar: 200 μm. **Right:** Localized magnification of the center. **d,** Projections and three-dimensional visualization of the pig brain. **Left:** Projections from different angles. Scale bar: 200 μm. **Center:** Three-dimensional visualization of the pig brain. Scale bar: 200 μm. **Right:** Localized magnification of the center.

We further evaluated the feasibility of acquiring large-scale imaging data through multiple overlapping scans in mouse brain imaging. During the two-dimensional imaging phase, 180 sagittal projections were acquired at 0°, followed by image registration (Supplementary Fig. 17). The registered images clearly delineated the sagittal plane, with high-magnification views revealing neuronal microstructures, including cell bodies and dendrites. For three-dimensional imaging, overlap rates were set to approximately 40% in the X direction and 20% in the Y direction. Figures 7b and 7c display two datasets along the X and Y axes, respectively, along with their registered three-dimensional reconstructions, which clearly depict neuronal cell bodies and dendrites with exceptional image quality. White arrows indicate the continuity of dendritic structures across datasets, highlighting the effectiveness of the imaging and spatial registration strategy. Additionally, registration accuracy in the Z direction was validated using a 20% overlap rate (Supplementary Fig. 18). This approach successfully acquired high-resolution imaging data from mouse cortical tissue, with a voxel size of 0.65 μm and a total volume of approximately 9 mm³. These results confirm the feasibility of this method for large-scale imaging and dataset registration (Supplementary Figs. 19 and 20, Supplementary Video 4).

**Fig. 7.**
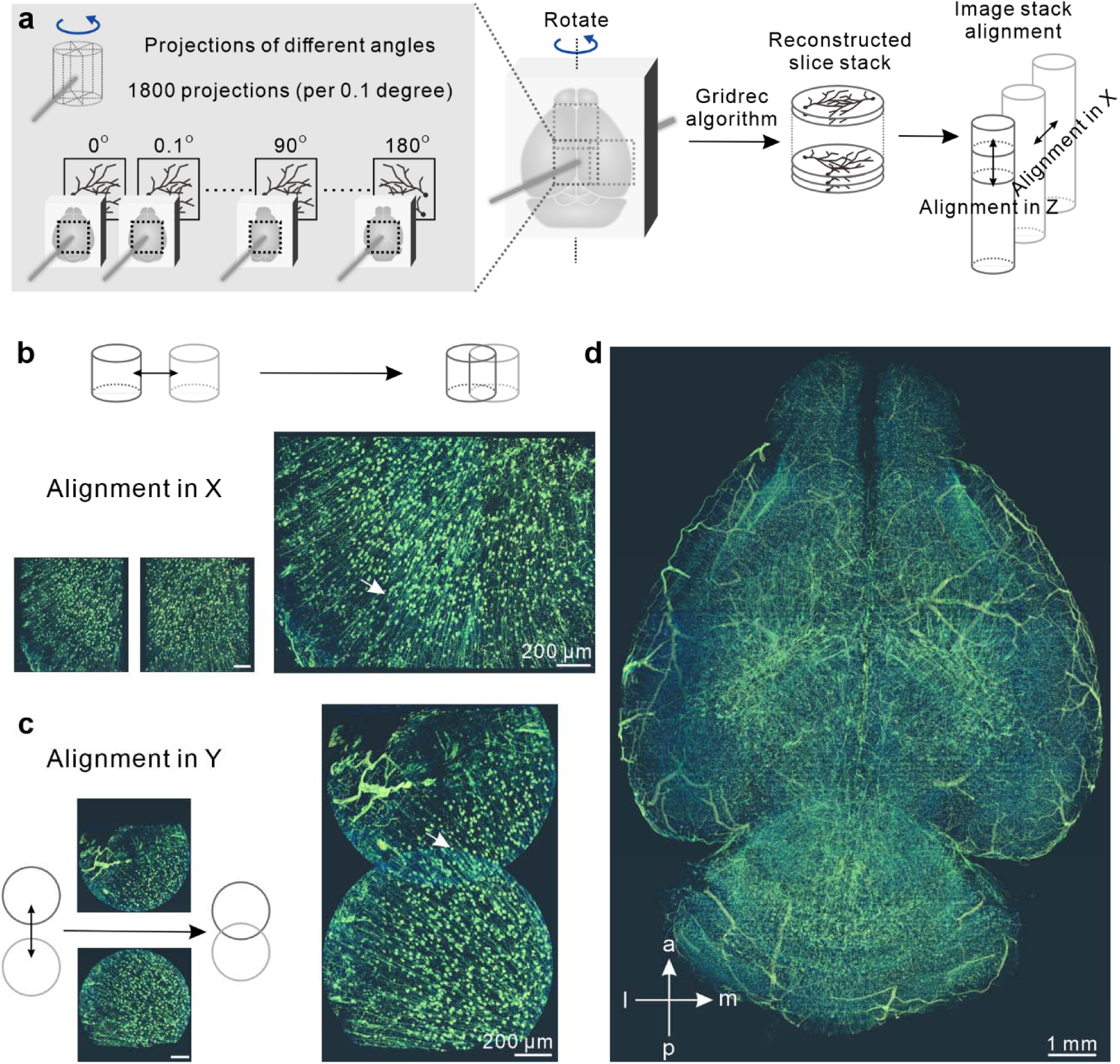
Synchrotron-based X-ray visualization of whole-brain neurons in a C57BL/6 mouse using HP-Golgi staining. **a,** Schematic of whole mouse brain X-ray imaging. In the horizontal direction, approximately 50% overlap was applied, and in the vertical direction, approximately 20% overlap was set. This configuration enabled the formation of a complete volume covering the entire mouse brain through multiple scans. For each Micro-CT scan, a total of 1,800 projection images were acquired over a 0 to 180-degree range. The Gridrec algorithm was then applied for slice reconstruction, followed by registration and visualization of the slices. **b,** Alignment of high-resolution mouse brain imaging data in the X-axis. **Top:** Schematic of X-axis alignment. **Bottom:** Registration of two image datasets (before and after alignment). Scale bar: 200 μm. **c,** Alignment of high-resolution mouse brain imaging data in the Y-axis. **Top:** Schematic of Y-axis alignment. **Bottom:** Registration of two image datasets (before and after alignment). Scale bar: 200 μm. **d,** Three-dimensional visualization of the whole mouse brain. Scale bar: 1 mm.

Finally, we acquired high-resolution Micro-CT imaging data of the intact mouse brain. Imaging was performed with a voxel size of 3.25 μm, and the field of view for each projection image measured approximately 6.7 mm × 3.3 mm. The anterior-posterior (AP) axis was designated as the rotation axis, enabling the acquisition of horizontal plane images at 0° and sagittal plane images at 90°. This configuration ensured that each CT scan fully captured the dorsal-ventral (DV) dimension of the brain, eliminating the need for registration along the DV axis. Whole-brain imaging was conducted at an X-ray energy of 25 keV. By seamlessly stitching together multiple Micro-CT scans, we successfully reconstructed a three-dimensional volume model of the entire mouse brain, with a total volume of approximately 1700 mm³. Figures 7d and Supplementary Video 5 present the three-dimensional visualizations, clearly revealing detailed microstructures such as neuronal cell bodies and cerebral vasculature. These results demonstrate the feasibility of combining the HP Golgi method with synchrotron-based X-ray microscopy for high-quality whole-brain imaging in mice. This approach establishes a robust foundation for future applications in large-scale brain imaging, including more complex and larger brain samples.

### Conclusions

In this study, we introduce the HP Golgi method, an innovative and reliable approach that substantially accelerates Golgi-Cox staining. By integrating high-pressure techniques, this method reduces the staining time for whole mouse (∼300 mm³) and rat (∼2000 mm³) brains to one-quarter of the original duration. For smaller zebrafish brains (∼0.03 mm³), staining time is reduced to 60% of the traditional method. Furthermore, we demonstrate rapid staining of hippocampal neurons in intact pig brains within 15 days, showcasing the enhanced staining speed for large brain tissue samples. Additionally, when coupled with synchrotron-based X-ray microscopy, the HP Golgi method enables high-resolution, three-dimensional imaging of the entire mouse brain.

The HP Golgi method offers several key advantages over existing neuron staining and labeling techniques. First, it ensures rapid penetration and staining. Unlike the classical Golgi-Cox method, which is hindered by the slow penetration of staining solutions, particularly when using fixatives like 4% paraformaldehyde, the HP Golgi method accelerates tissue infiltration via high-pressure technology. This improvement not only reduces staining time but also addresses the classical method’s limitations in staining deeper neurons, which often results in tissue autolysis. In contrast, the HP Golgi method provides more effective internal neuron staining and fixation within a shorter time frame, preserving structural integrity and minimizing tissue degradation. Second, the HP Golgi method is ethically more applicable for ex vivo human brain research. While viral-based neuron labeling techniques face significant ethical concerns, the HP Golgi method circumvents these issues, making it suitable for use with human brain tissue ^36^. This opens new avenues for investigating human neuronal structures and functions. However, our experiments with whole porcine brains (∼10□ mm³) reveal a limitation. Despite the accelerated penetration in smaller regions, deeper areas remain susceptible to tissue autolysis due to insufficient staining solution diffusion. This issue persists, hindering clear staining and imaging of cortical neurons in large porcine brains. Future work will focus on addressing these challenges, particularly in large primate and human brain tissue, by further optimizing the HP Golgi method and integrating additional technologies to prevent tissue degradation over time. Finally, the HP Golgi method offers superior imaging capabilities. The staining solution contains mercury (Hg) and chromium (Cr), both of which exhibit strong hard X-ray absorption, enabling clear visualization of neuronal structures through Micro-CT imaging. Recent advances in synchrotron-based X-ray holographic nano-tomography (XNH) ^37^ and nano-computed tomography (Nano-CT) have enabled imaging resolutions approaching those of electron microscopy. As these technologies continue to improve ^38–41^, the HP Golgi method holds significant potential for nanoscale, three-dimensional imaging of neuronal networks, particularly in studies of complex brain structures and functional connectivity.

## Methods

### Animals

Mice: C57BL/6 mice (6-8-week-old, male) were purchased from Shanghai SLAC Laboratory Animal Co., Ltd., China. Zebrafish: AB Zebrafish (12-month-old, female) were purchased from Shanghai FishBio Co., Ltd., China. Rats: 6-week-old SD rats (6-week-old, male) were purchased from Shanghai SLAC Laboratory Animal Co., Ltd., China. Pigs: Bama pigs (2-month-old, male) were purchased from Hangzhou tuoshi Biotechnology Co., Ltd., China. All the animal experiments were conducted in accordance with the Institute’s Guide for the Care and Use of Laboratory Animals and were approved by the ethical committee of Shanghai Beautiful Life Medica Technology (approval no. SYXK-2017-0016, approved on 25 December 2017).

### Golgi-Cox method and HP Golgi method

Before taking the brain, all model animals were perfused with normal saline through the heart. Brains were quickly removed from the skull and washed with normal saline water to remove blood from the surface. Golgi impregnation of neurons was performed using the Hito Golgi-Cox OptimStain^TM^ Kit. For Golgi-Cox method, samples were dyed at 26 and atmospheric pressure in the dark. For HP Golgi method, samples were dyed at 37 and 9 MPa in the dark. The brain samples were placed in sealed bags containing Golgi staining solution, and any residual air was carefully expelled. The samples were subsequently placed in a high-pressure assisted device. The rates of pressurization and depressurization were deliberately slowed to prevent rapid pressure changes from damaging the brain samples. Throughout the staining process, a high-pressure environment and a temperature of 37°C were maintained.

### Micro-CT Experiment

Micro-CT imaging data were acquired at beamlines of the synchrotron radiation source. The two-dimensional imaging data were collected at the TPS 02A1 brain imaging beamline of Taiwan Photon Source (TPS, China), while the three-dimensional imaging data were obtained from the BL13HB X-ray imaging and biomedical application beamline and BL16U2 fast X-ray imaging beamline of the Shanghai Synchrotron Radiation Facility (SSRF, China). For the two-dimensional imaging data, the pixel size was set to 0.5 μm, with the imaging energy of 14 keV. Two-dimensional projections with both lateral and longitudinal overlap were acquired, and the panoramic images of the brain slice were ultimately stitched together using the Stitching – Grid/Collection Stitching tool in Fiji ImageJ. The voxel size for the three-dimensional imaging data was 0.65 μm, with the imaging energy similarly set at 14 keV. Projections were acquired at equal intervals within a range of 0 to 180°, resulting in either 900 or 1800 projections.

### Whole Mouse Brain Imaging

The Micro-CT imaging data of whole mouse brain were obtained from the BL13HB X-ray imaging and biomedical application beamline. The pixel size was set to 3.25 μm, with the imaging energy of 25 keV. Projections were acquired at equal intervals within a range of 0 to 180°, resulting in 1800 projections. The anterior-posterior (AP) axis of the mouse brain was designated as the rotation axis, allowing us to obtain horizontal plane image of the entire brain at 0° and sagittal plane image at 90°. In the medial-lateral (LM) axis of the mouse brain, we established an overlap rate of approximately 50% by using four overlapping projection images to achieve comprehensive coverage of the entire mouse brain. For the anterior-posterior (AP) axis, the overlap rate was set at around 20%, utilizing five overlapping projection images to ensure full coverage. In the dorsal-ventral (DV) axis, a single set of data was sufficient to cover the entire mouse brain. Ultimately, we constructed a three-dimensional volume that encompasses the entire mouse brain by stitching together 20 sets of data.

### Reconstruction and Three-Dimensional Visualization

The projections were reconstructed into slices using the PITRE software, employing the Gridrec algorithm. The reconstructed slices were processed using Fiji ImageJ software to remove circular boundaries. Three-dimensional visualization was performed using Amira software. For the three-dimensional visualization of the entire mouse brain, data from different groups were imported into Amira, where the Register Images and Merge modules were utilized for data registration and fusion, respectively.

## Data availability

The data that support the findings of this study are available within the paper and its Supplementary Information files. Source data are provided with this paper.

## Acknowledgements

We thank the staff of BL13HB X-ray imaging and biomedical application beamline and BL16U2 fast X-ray imaging beamline of Shanghai Synchrotron Radiation Facility (SSRF, China), and TPS 02A brain imaging beamline of Taiwan Photon Source (TPS, China) for their technical support and assistance. This work was supported by the National Key R&D Program of China (2022YFA1603600 to J.H.), the National Natural Science Foundation of China (T2188102 to C.F., 21991134 to C.F., and 22325406 to J.L.), 2022 Shanghai “Science and Technology Innovation Action Plan” Fundamental Research Project (22JC1401203 to Y.Z.).

## Author contributions

C.F. and J.H. supervised the research. Q.T., X.C., Y.L. and Y.Z. performed the experiments. X.Y. and F.Z. helped in guidance of animal experiments. J.Z., K.L. and B.D. helped in guidance of X-ray microscopy. Q.L. and L.W. performed critical revisions. J.L., Y.Z. and C.F. designed the experiments, analyzed the data and wrote versions of manuscript. All authors discussed and commented on the manuscript.

## Competing interests

The authors declare no competing interests.

## Notes

### Competing Interest Statement

The authors have declared no competing interest.

