## Supporting Information for "High-pressure Golgi neuronal staining for X-ray whole-brain imaging"

**Supplementary methods**

**Tissue Protection, Sectioning, Developing and Sealing Step.**

For each brain sample, it is transferred from Golgi staining solution to tissue-protectant solution (30% sucrose solution) and stored at 4°C in dark. After every 24 h, each brain sample is transferred into a new small bottle with fresh tissue-protectant solution. The samples are kept at 4°C in dark for 3 days. For frozen microtome sectioning, brains were embedded in Tissue Tek O.C.T. compound, cryo-sectioned sagittal at 100 mm thickness and directly mounted on gelatin-coated slides with the help of solution 3 provided in the kit. Subsequently, the sections were developed in the mixed solution of solution 4 and solution 5, and then dehydrated by gradient ethanol, and cleared with xylene, and finally sealed with the help of Eukitt.

**Optica Microscopy.**

Brain slices from different model organisms were imaged using ZEISS Axio Imager 2 optical microscope. Under the same imaging parameters, images of the regions of interest (ROI) and background were captured, with background removal performed using Fiji ImageJ software. Panoramic images were also acquired on the ZEISS Axio Imager 2 optical microscope, with both horizontal and vertical repeat rates set, and background images captured under the same imaging parameters, followed by background removal using Fiji ImageJ software. Finally, the panoramic images of the brain slices were stitched together using the Stitching - Grid/Collection Stitching tool in Fiji ImageJ.

**Sholl Analysis.**

To detect the effect of the HP Golgi method on the staining of neuronal structures, we conducted Sholl analysis on cortical neurons from various model organisms. For each dataset, we selected at least 10 neurons for statistical analysis. Stained neurons were imaged with the ZEISS Axio Imager 2 optical microscope under 40x or 20x objective lens. Manual tools were used to trace each branch of the neurons. Subsequently, the number of intersections was calculated using the Sholl analysis tool in Fiji ImageJ, and the data were processed using Origin software.

**Sample Preparation for Micro-CT.**

The brain samples of different animal models that had been processed with a 30% sucrose solution were developed in mixed solutions 4 and 5 for a staining procedure, which lasted for 6 hours with solution changes every 2 hours. Subsequently, the samples were dehydrated using a gradient of ethanol, culminating in an overnight treatment in 100% ethanol. The brain samples were then transferred to a solution composed of a 1:1 mixture of ethanol and xylene, where they were immersed at 40°C for 1 hour. Following this, the samples underwent a clearing process in xylene, repeated three times for 1 hour each, while maintaining a temperature of 40°C. Finally, the brain tissues were transferred to molten paraffin for 2 hours before being embedded using a paraffin embedding machine.

**Supplementary figures.**

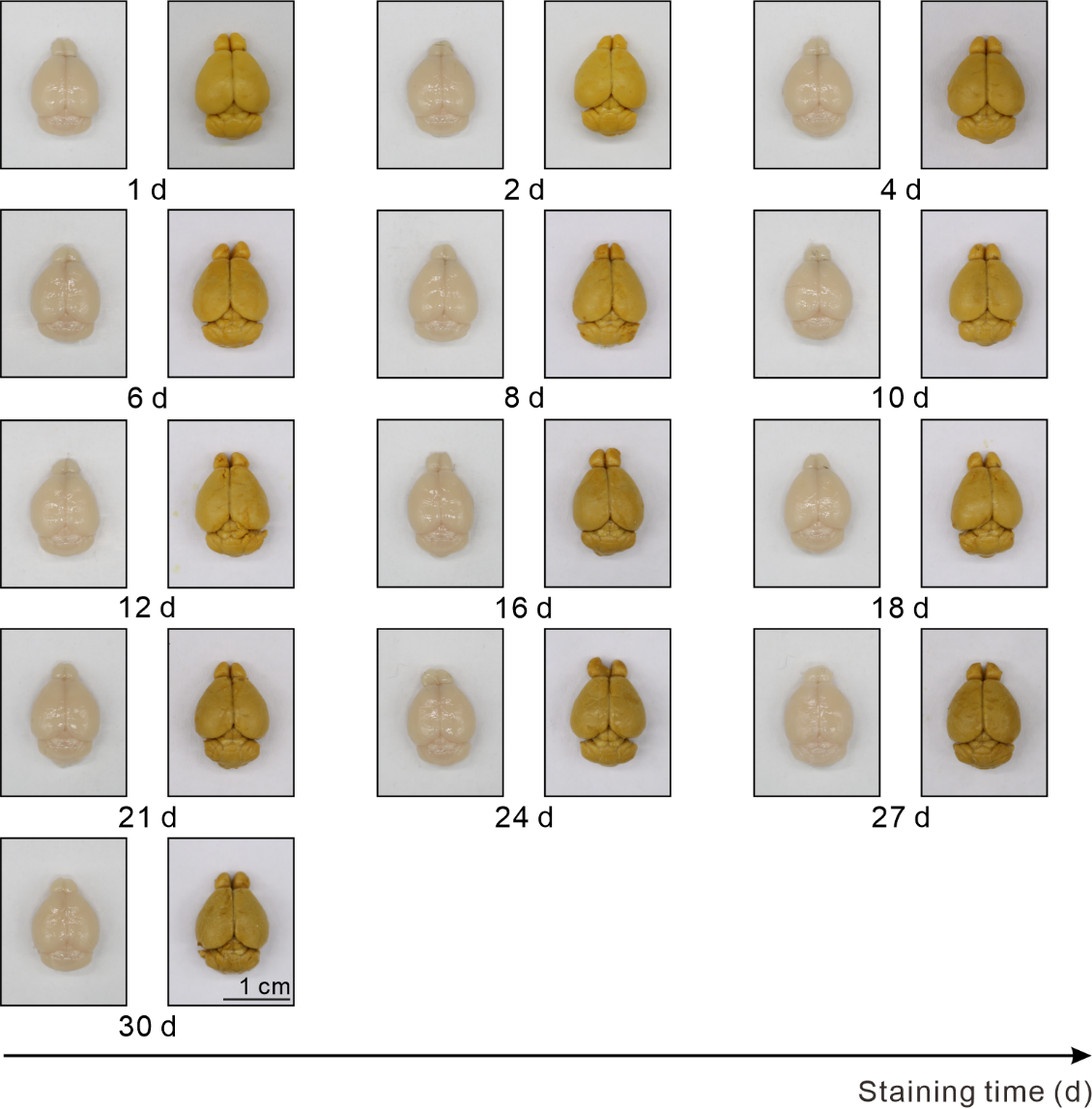

**Supplementary Figure 1. Images of mouse brains stained at different time points using the Golgi-Cox method, viewed from the front and back.** As the staining time increased, the color of the mouse brains transitioned from bright yellow to brownish yellow. Scale bar: 1 cm.

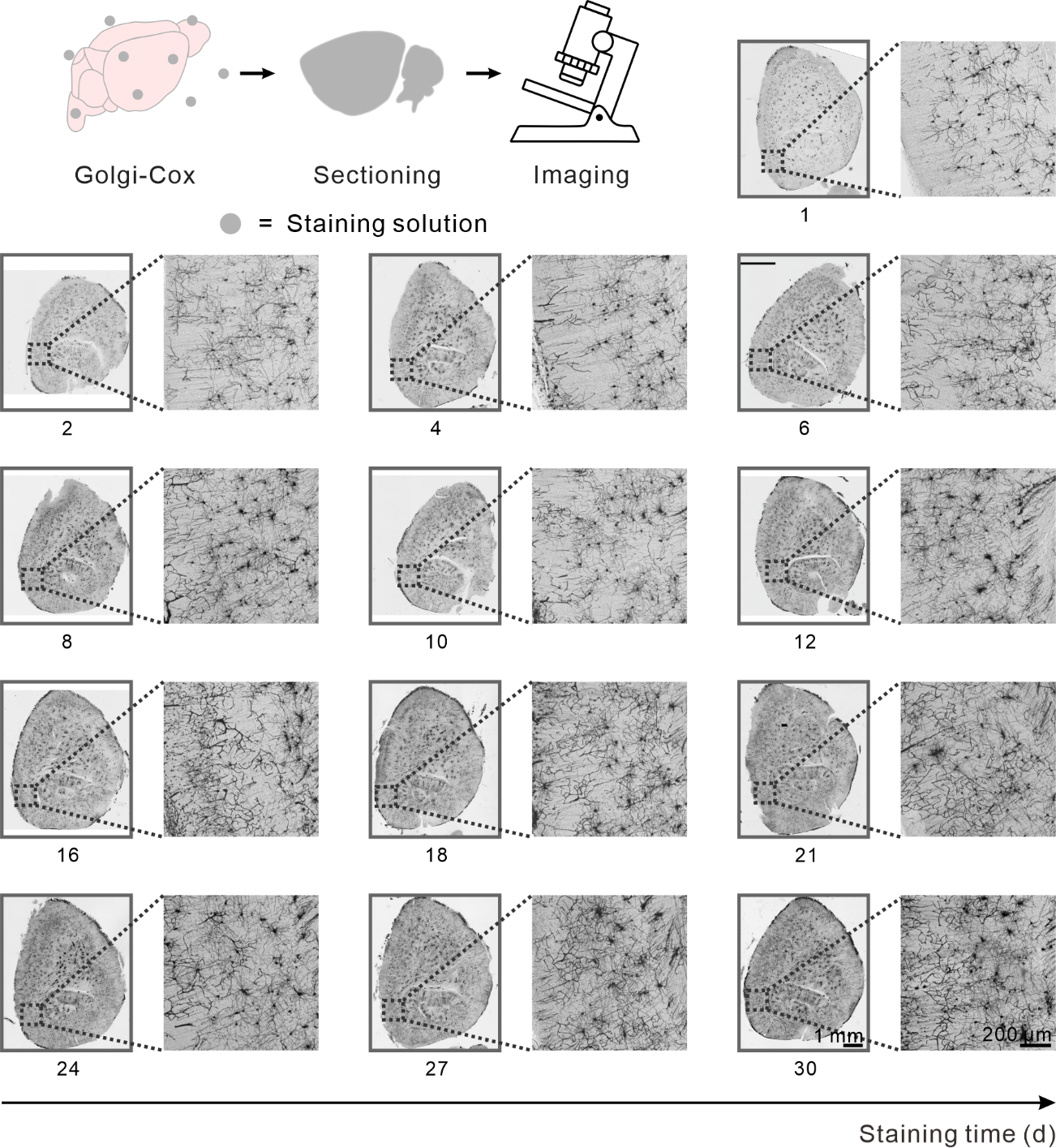

**Supplementary Figure 2. Panoramic and magnified images of mouse brain sections stained using the Golgi-Cox method for varying durations.** For staining periods between 1 and 16 days, neuronal morphology was clearly discernible, with minimal blood vessel staining. However, from 18 to 30 days, a significant increase in stained blood vessels was observed. Scale bars: 1 mm for the panoramic image and 200 μm for the magnified view.

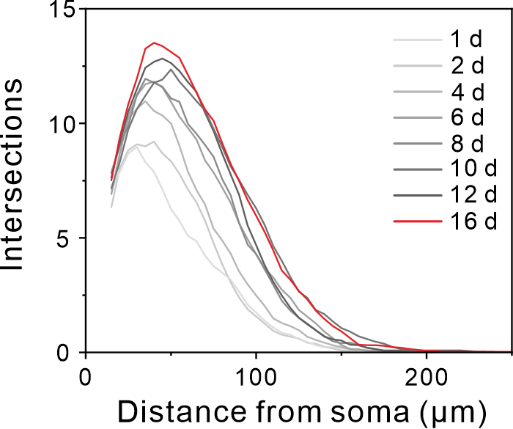

**Supplementary Figure 3. Sholl analysis of somatosensory (SS) neurons stained for different durations using the Golgi-Cox method.** Longer staining times resulted in an increased number of intersections at proximal distances (20–60 μm) and extended dendritic branches at distal locations (>150 μm), indicating progressively more complete visualization of neuronal structures.

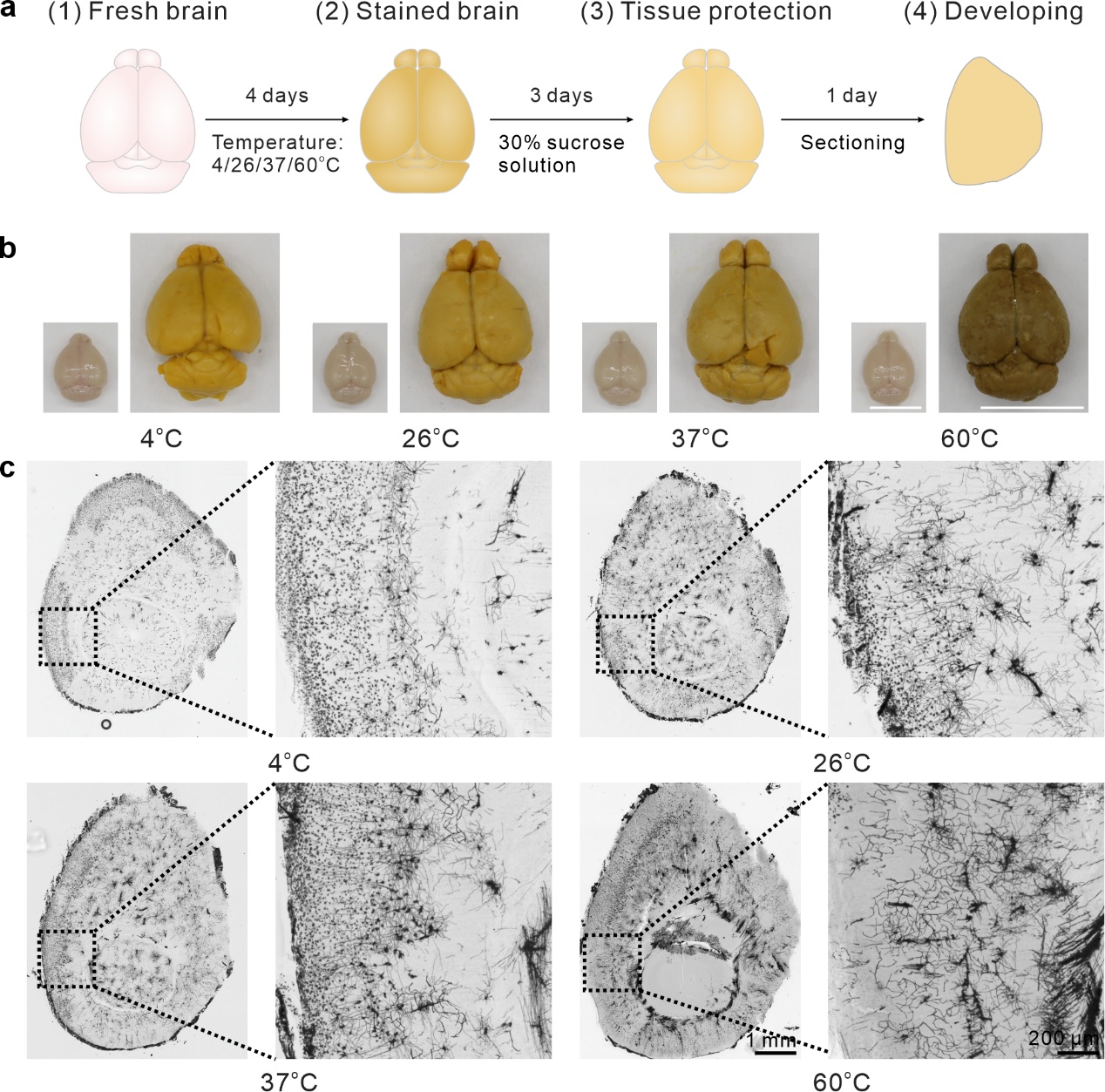

**Supplementary Figure 4. Effects of different environmental temperatures on mouse brain staining. a,** Schematic workflow for the staining, sectioning, and imaging processes. **b,** Images of mouse brains before and after staining. As the environmental temperature increases, the color of the mouse brains transitions from bright yellow to dark brown. Scale bar: 1 cm. **c,** Panoramic and magnified images of stained mouse brain sections. At 4°C, 26°C, and 37°C, neuronal structures are distinctly visible in brain slices, with minimal staining of blood vessels. However, at 60°C, neuronal structures become challenging to observe, accompanied by a substantial increase in stained blood vessels. Scale bar: 1 mm for panoramic images and 200 μm for magnified views.

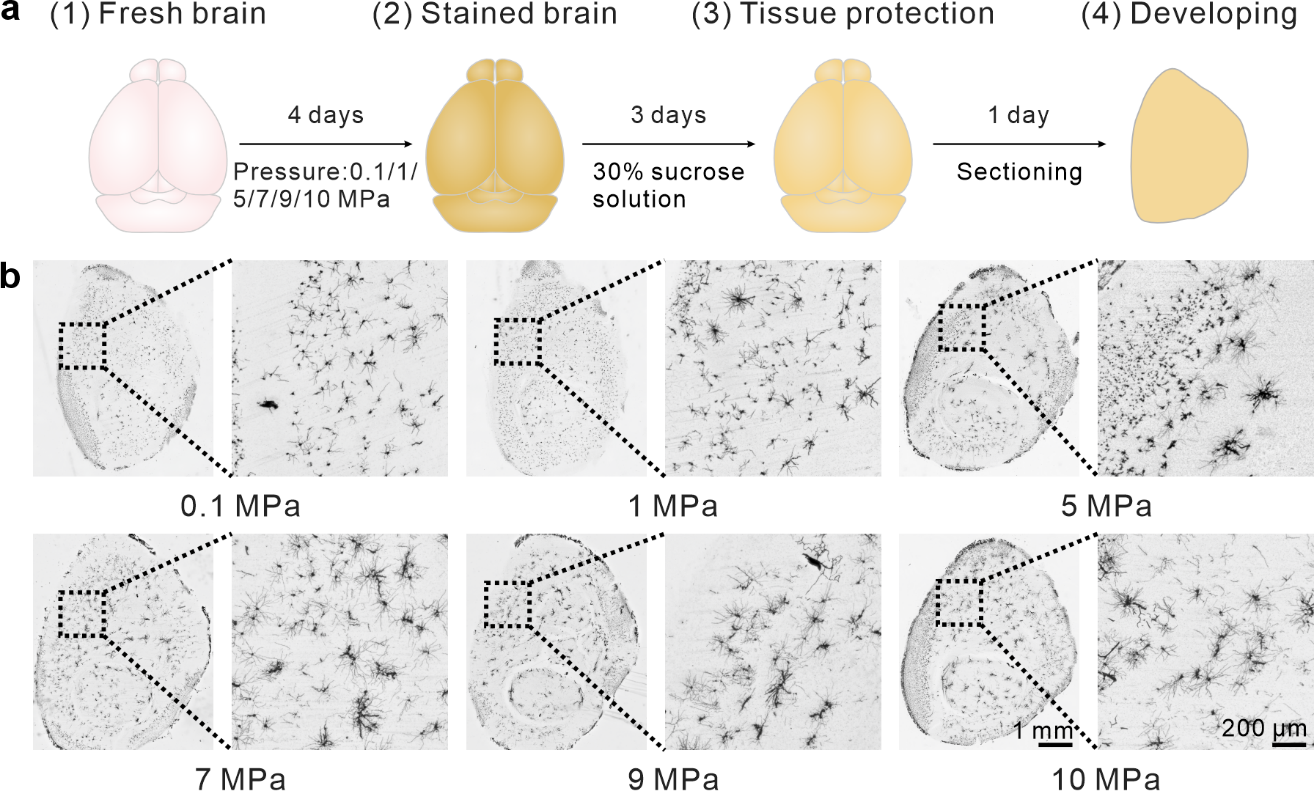

**Supplementary Figure 5. Effects of different pressures on mouse brain staining. a,** Schematic workflow for the staining, sectioning, and imaging processes. **b,** Panoramic and magnified images of stained mouse brain sections. Panoramic images show no significant tissue damage across different pressures, with a relatively low background signal. Magnified images of the SS region reveal well-defined neuronal morphologies at all pressures, with minimal vascular staining. Scale bar: 1 mm for panoramic images and 200 μm for magnified views.

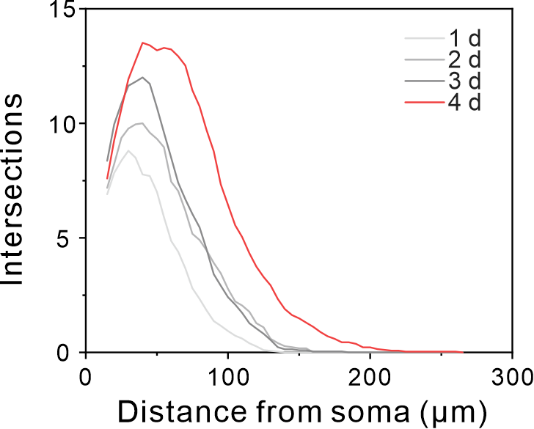

**Supplementary Figure 6. Sholl analysis of SS neurons stained for varying durations using the HP Golgi method.** As staining time increased, neurons exhibited a greater number of intersections at proximal distances (20-60 μm) and extended dendritic branches at distal locations (>150 μm), indicating that neuronal structures became increasingly more complex and complete over time.

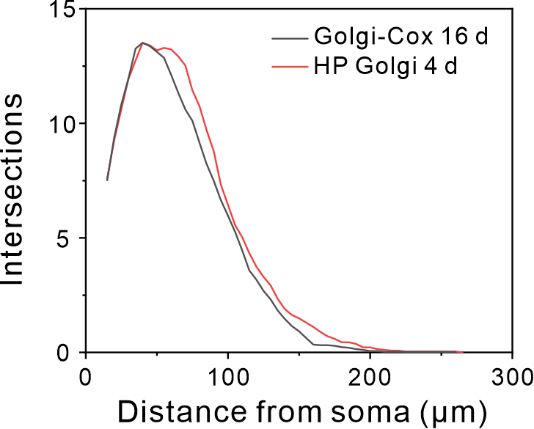

**Supplementary Figure 7. Comparison of the HP Golgi method and the Golgi-Cox method.** Mouse brains were stained for 4 days using the HP Golgi method and for 16 days using the Golgi-Cox method. Sholl analysis of neurons in the SS brain region from both staining methods showed no significant differences in neuronal complexity.

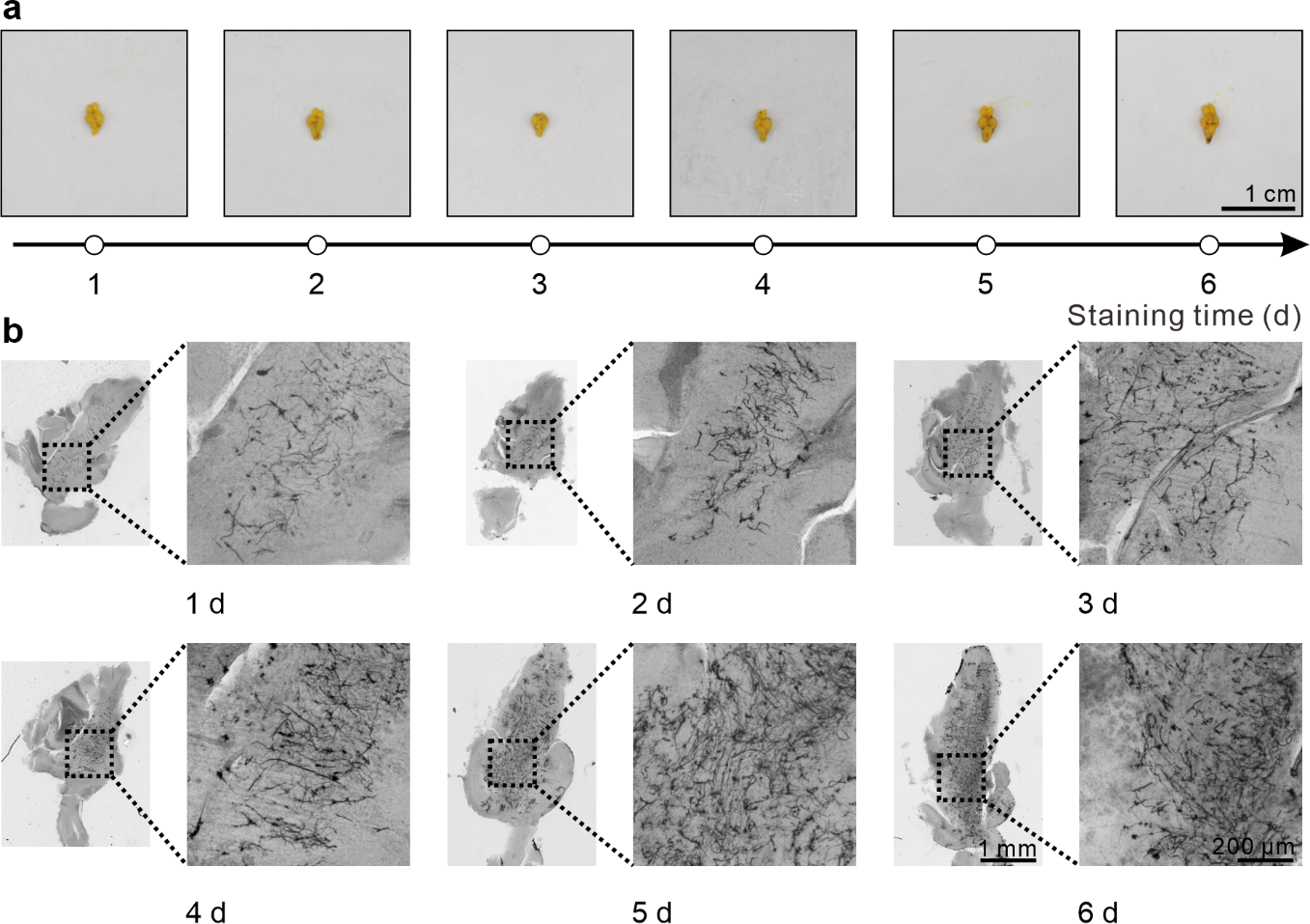

**Supplementary Figure 8. Results of zebrafish brains stained for varying durations using the Golgi-Cox method. a,** Images of zebrafish brains post-staining at 1, 2, 3, 4, 5, and 6 days. Scale bar: 1 cm. **b,** Panoramic images and magnified views of zebrafish brain sections. Neuronal morphology was clearly discernible across all time points. However, by day 6, increased vascular staining compromised the clarity of neuronal observations. Scale bar: 1 mm for panoramic images and 200 μm for magnifications.

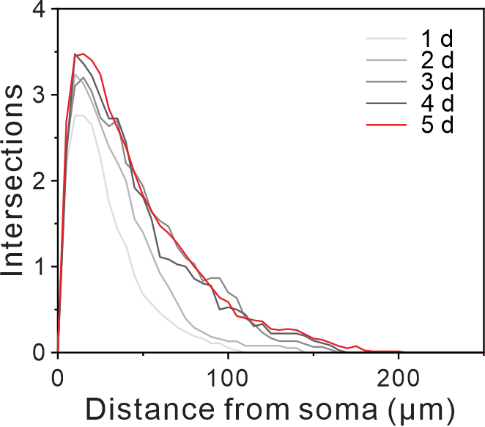

**Supplementary Figure 9. Sholl analysis of zebrafish brain neurons stained for varying durations using the Golgi-Cox method.** With longer staining times, neuronal morphology exhibited progressive completeness, as evidenced by an increase in intersections and extended dendritic branches.

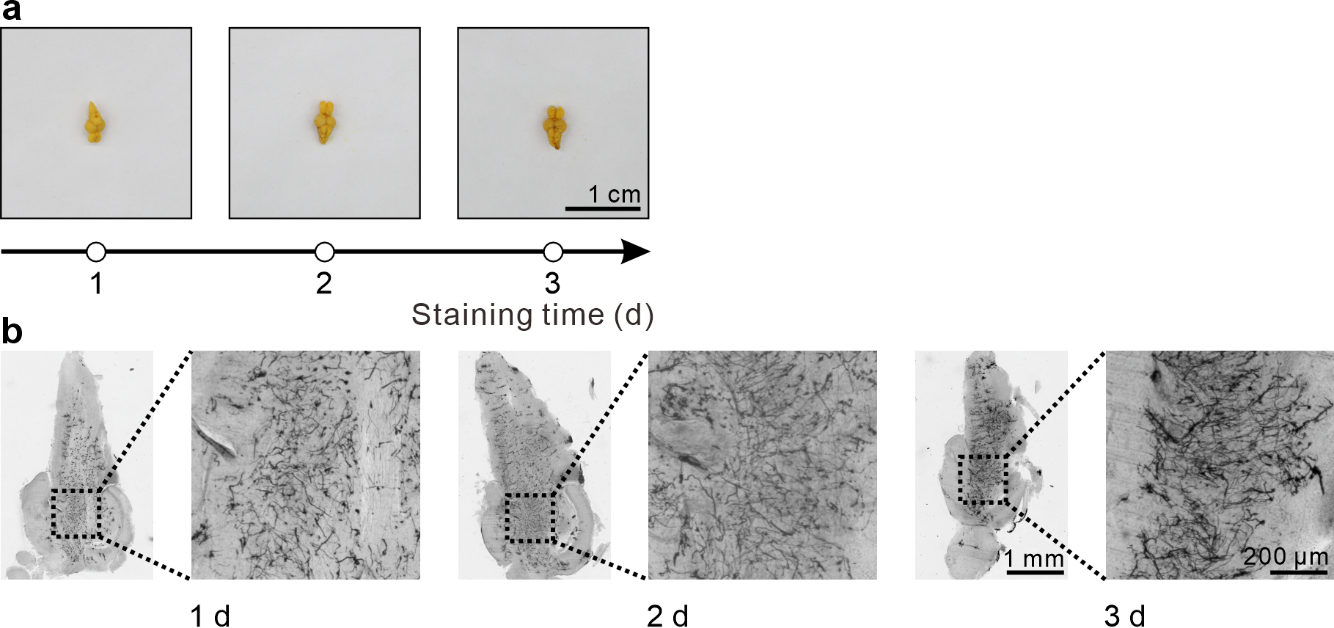

**Supplementary Figure 10. Results of zebrafish brains stained for different durations using the HP Golgi method. a,** Images of zebrafish brains after staining for 1, 2, and 3 days. Scale bar: 1 cm. **b,** Panoramic images and magnified views of zebrafish brain sections. Distinct neuronal morphology was consistently observed across all time points. Scale bar: 1 mm for panoramic images and 200 μm for magnified views.

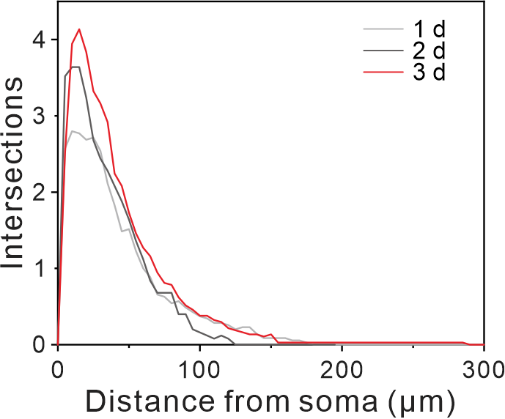

**Supplementary Figure 11. Sholl analysis of zebrafish brain neurons stained for different durations using the HP Golgi method.** Neurons stained for 3 days exhibited more complete and complex structures compared to those stained for 1 or 2 days.

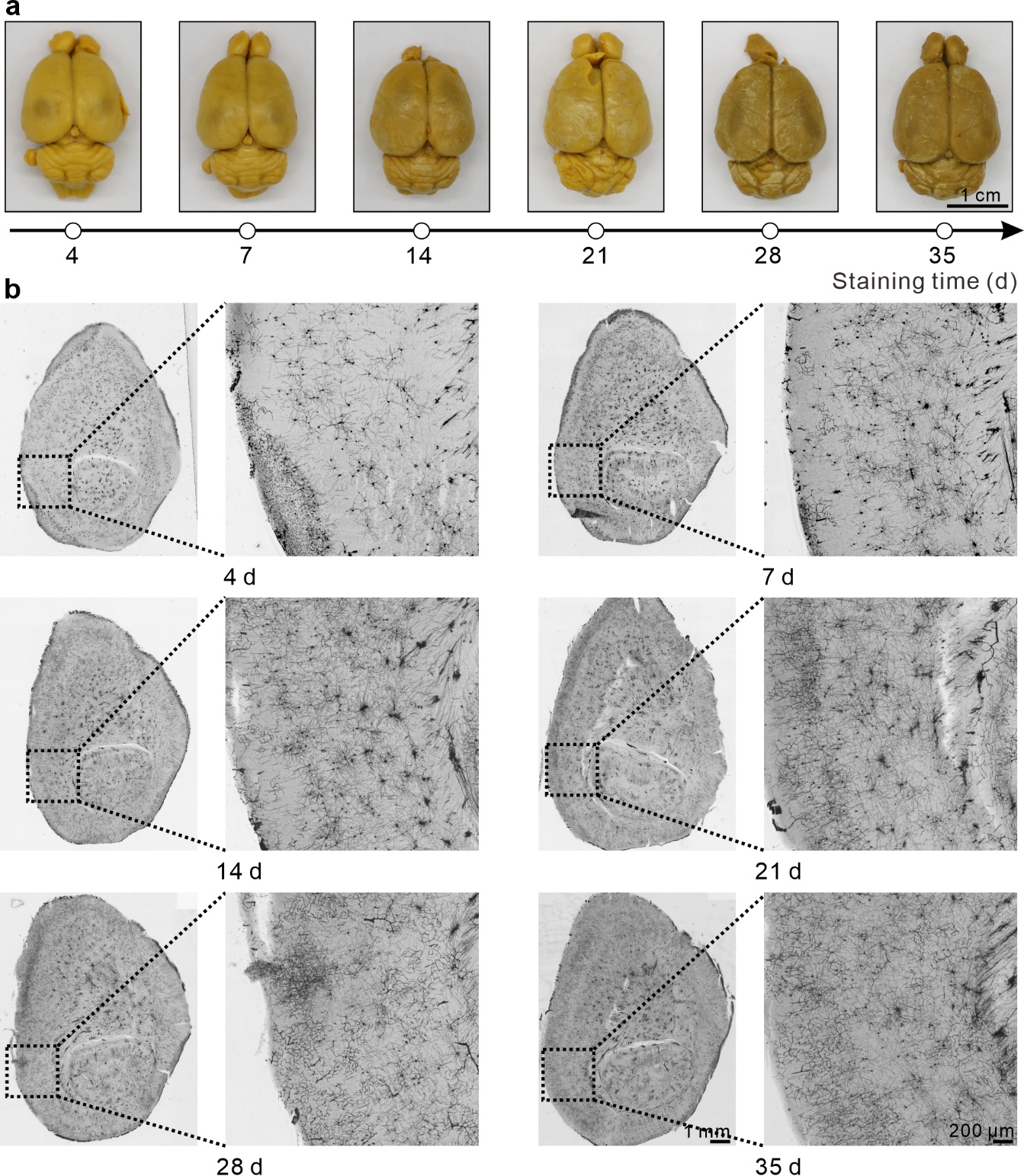

**Supplementary Figure 12. Results of rat brains stained for different durations using the Golgi-Cox method. a,** Images of rat brains after staining for 4, 7, 14, 21, 28, and 35 days. The brain color transitioned from bright yellow to darker tones with longer staining durations. Scale bar: 1 cm. **b,** Panoramic and magnified images of rat brain sections. Neuronal morphology was clearly observable at staining durations between 4 and 28 days, accompanied by minor vascular staining. At 35 days, vascular staining significantly increased, interfering with neuronal visualization. Scale bar: 1 mm for panoramic images and 200 μm for magnified images.

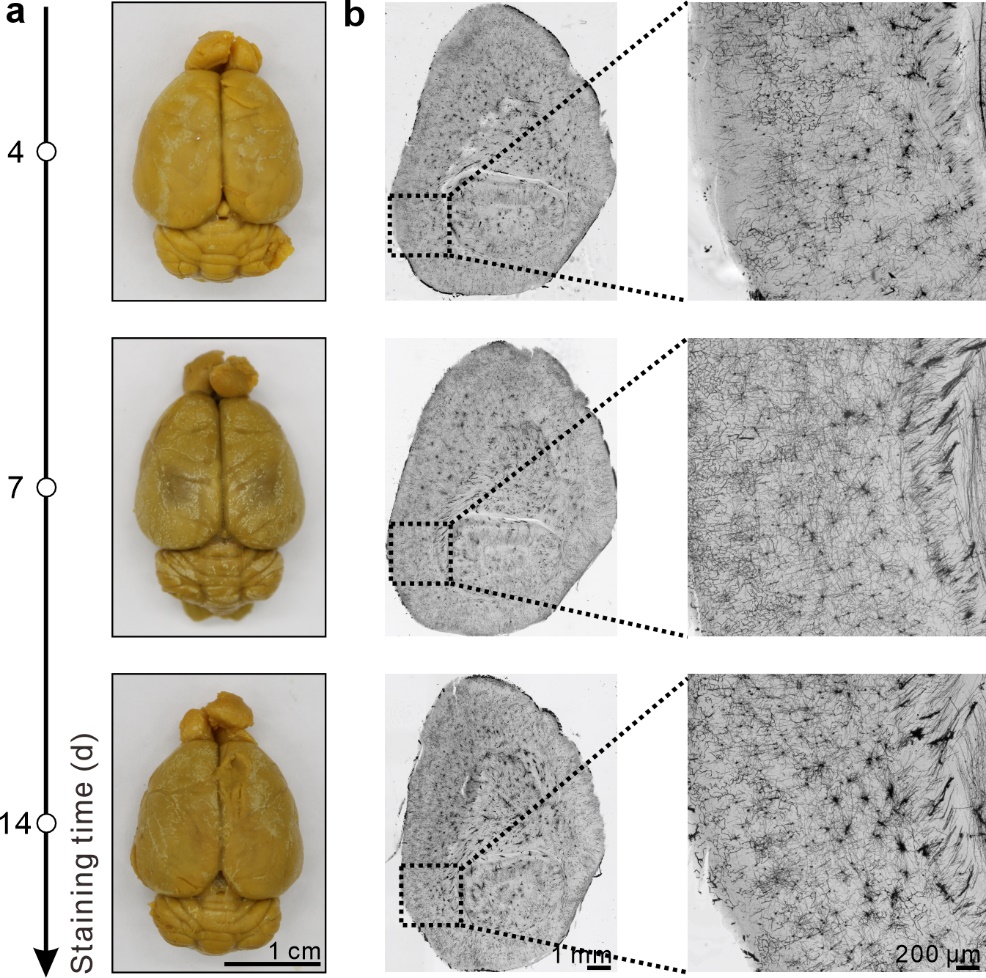

**Supplementary Figure 13. Results of rat brains stained for different durations using the HP Golgi method. a,** Images of rat brains after staining for 4, 7, and 14 days. The brain color progressively darkened with longer staining durations. Scale bar: 1 cm. **b,** Panoramic and magnified images of rat brain sections. Neuronal morphology was clearly observed at all three time points. However, an increase in the number of stained blood vessels was noted at 14 days, which could interfere with neuronal visualization. Scale bar: 1 mm for panoramic images and 200 μm for magnified images.

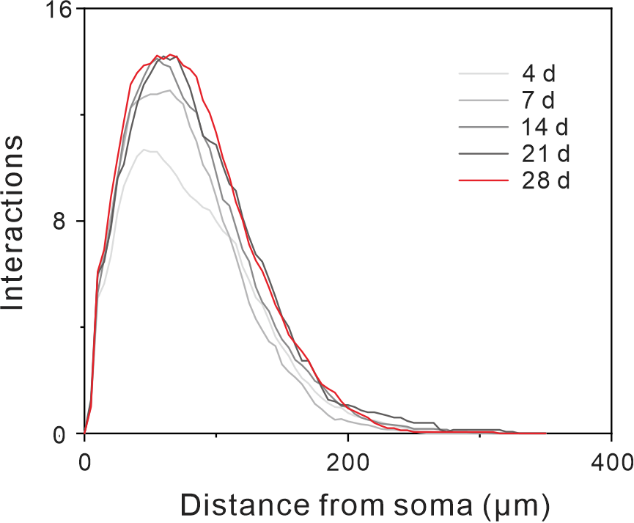

**Supplementary Figure 14. Sholl analysis of rat brain neurons stained for varying durations using the Golgi-Cox method.** Prolonged staining times resulted in progressively more complete neuronal morphology in the SS region, as indicated by an increase in the number of proximal intersections (20–60 μm) and extended dendritic branches at distal locations (>150 μm).

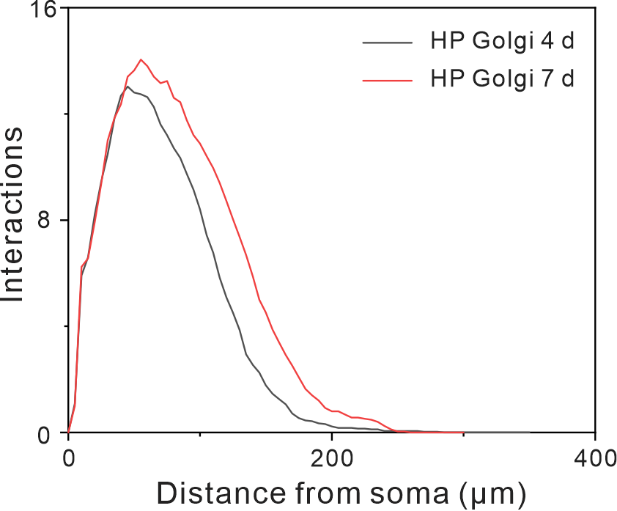

**Supplementary Figure 15. Sholl analysis of rat brain neurons stained for varying durations using the HP Golgi method.** Longer staining times resulted in progressively more complete neuronal morphology in the SS region, reflected by an increased number of intersections at proximal distances (20–60 μm) and extended dendritic branches at distal locations (>150 μm).

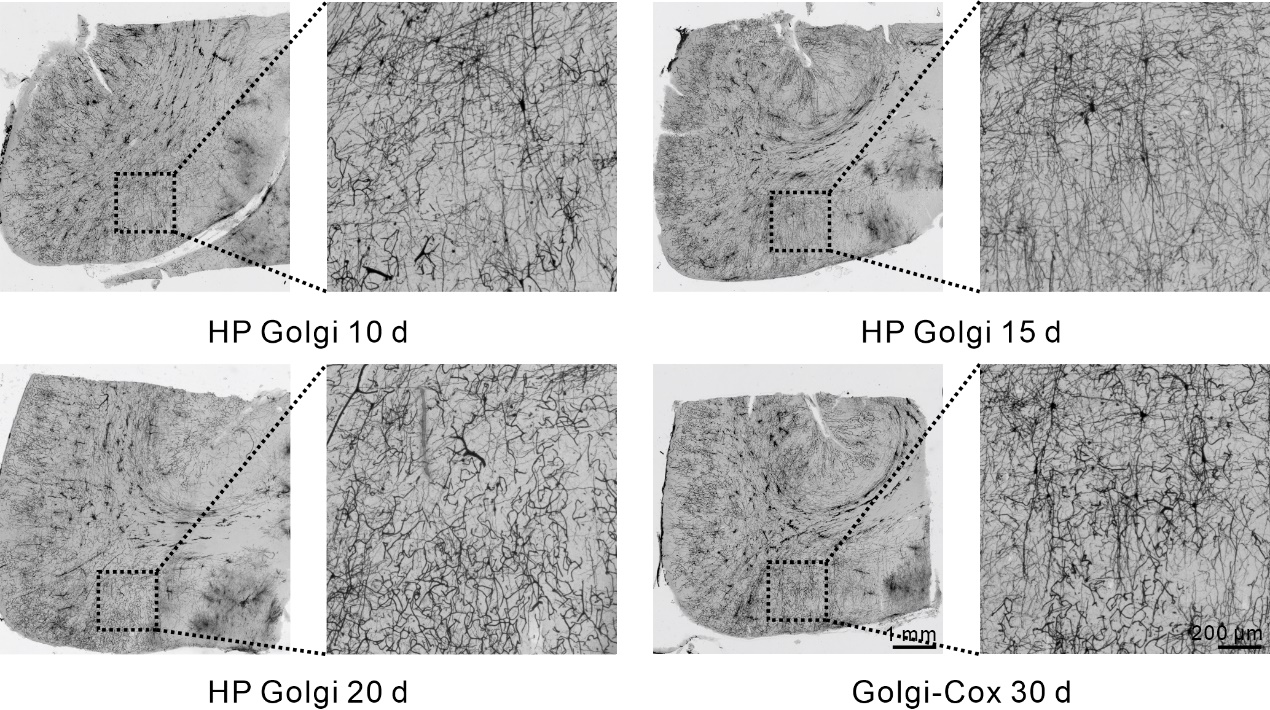

**Supplementary Figure 16. Comparative analysis of rat brains stained for different durations.** The Golgi-Cox method was applied for 30 days, while the HP Golgi method was used for 10, 15, and 20 days. Neuronal structures became progressively more complete with longer staining times using the HP Golgi method; however, at 20 days, prominent vascular staining was observed. Notably, the 15-day HP Golgi staining group showed reduced vascular staining and superior visualization of hippocampal neurons compared to the 30-day Golgi-Cox method. Scale bar: 1 mm for panoramic images and 200 μm for magnified images.

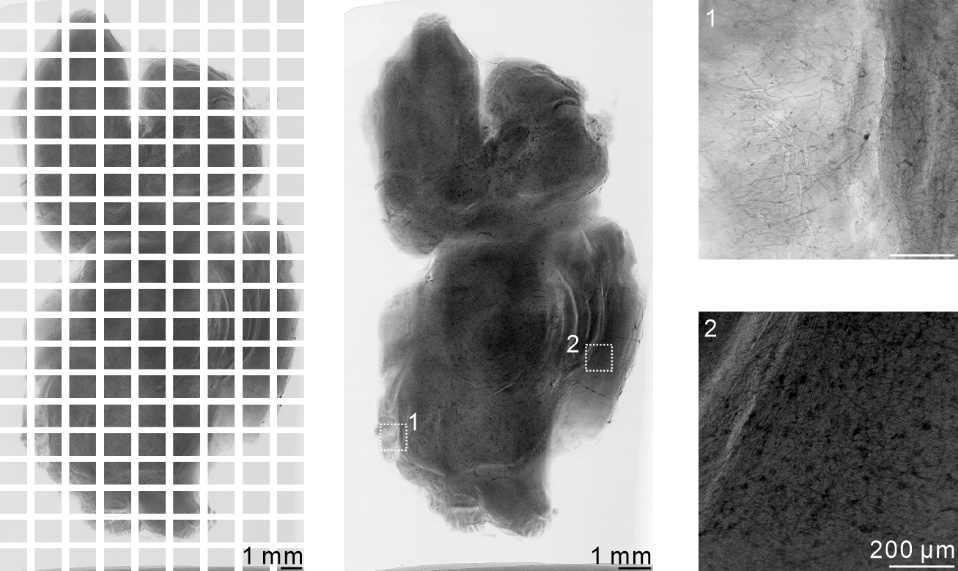

**Supplementary Figure 17. Two-dimensional stitching of the sagittal brain section in mice.** Using a total of 180 two-dimensional projection images—organized into 9 horizontal and 20 vertical groups—sagittal sections of the mouse brain were successfully reconstructed through image registration. Scale bar: 1 mm for panoramic images and 200 μm for magnified images.

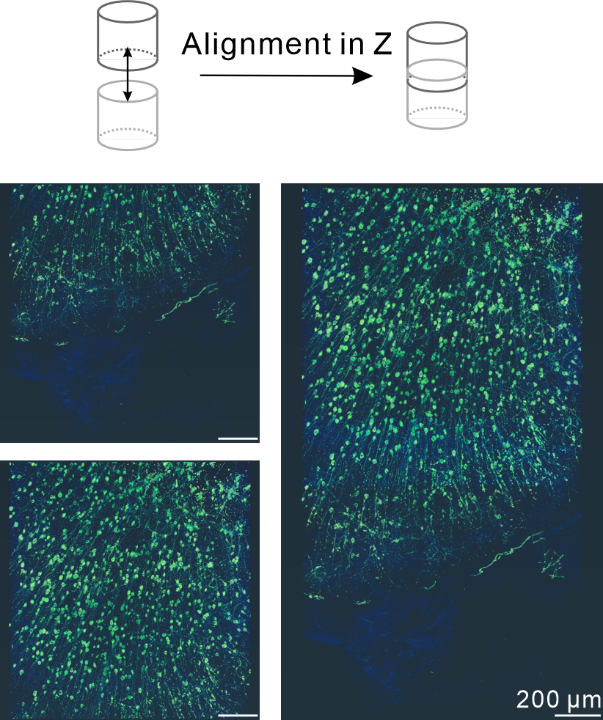

**Supplementary Figure 18. Alignment of high-resolution mouse brain imaging data in the Z-axis. a,** Schematic representation of the Z-axis alignment process. **b,** Registration of two sets of image data, showing the alignment of before and after images. Scale bar: 200 μm.

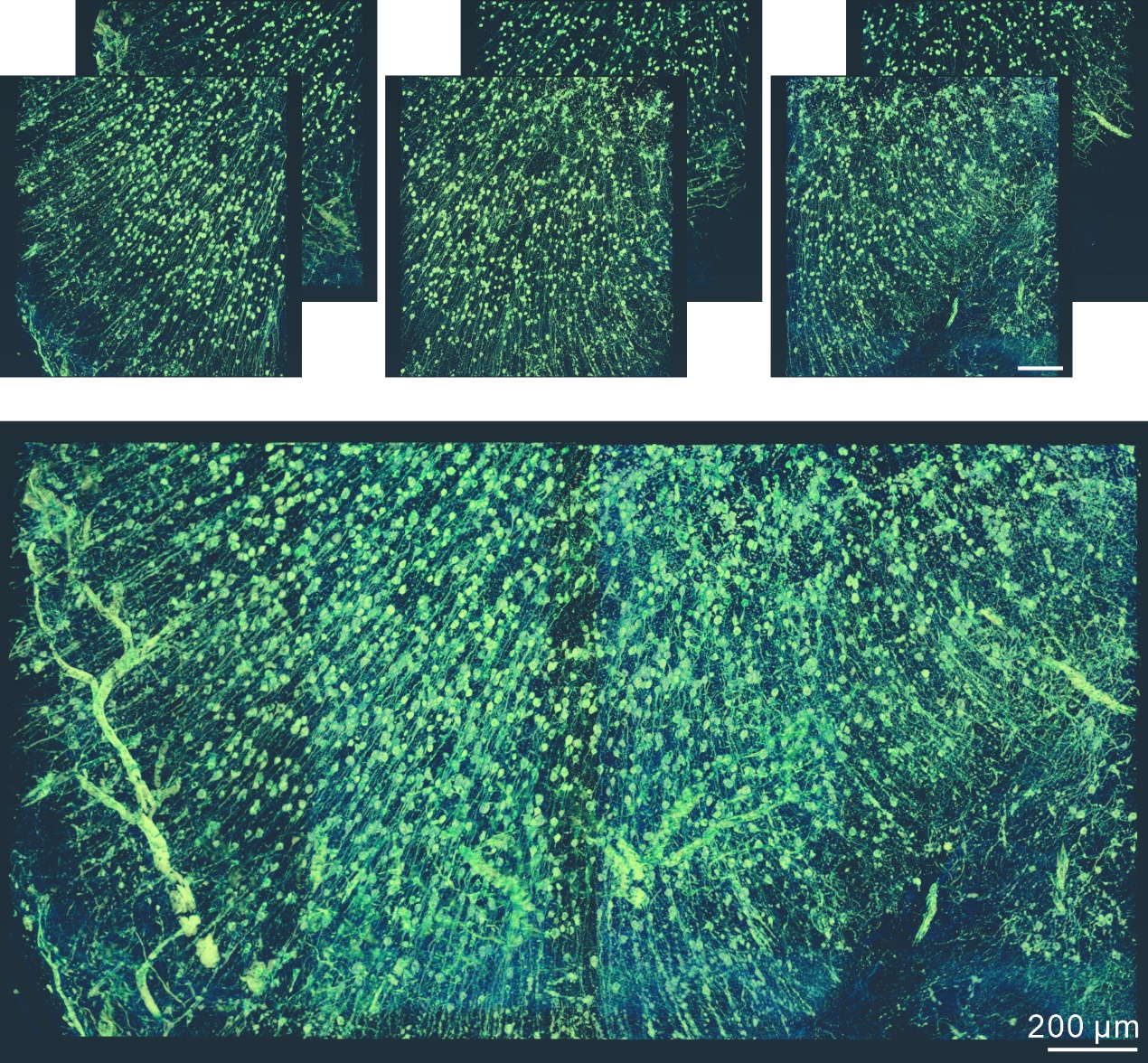

**Supplementary Figure 19. Three-dimensional visualization of the mouse brain cortex (XZ plane).** A 3D volume of approximately 9 mm³ was generated by combining six datasets. Scale bar: 200 μm.

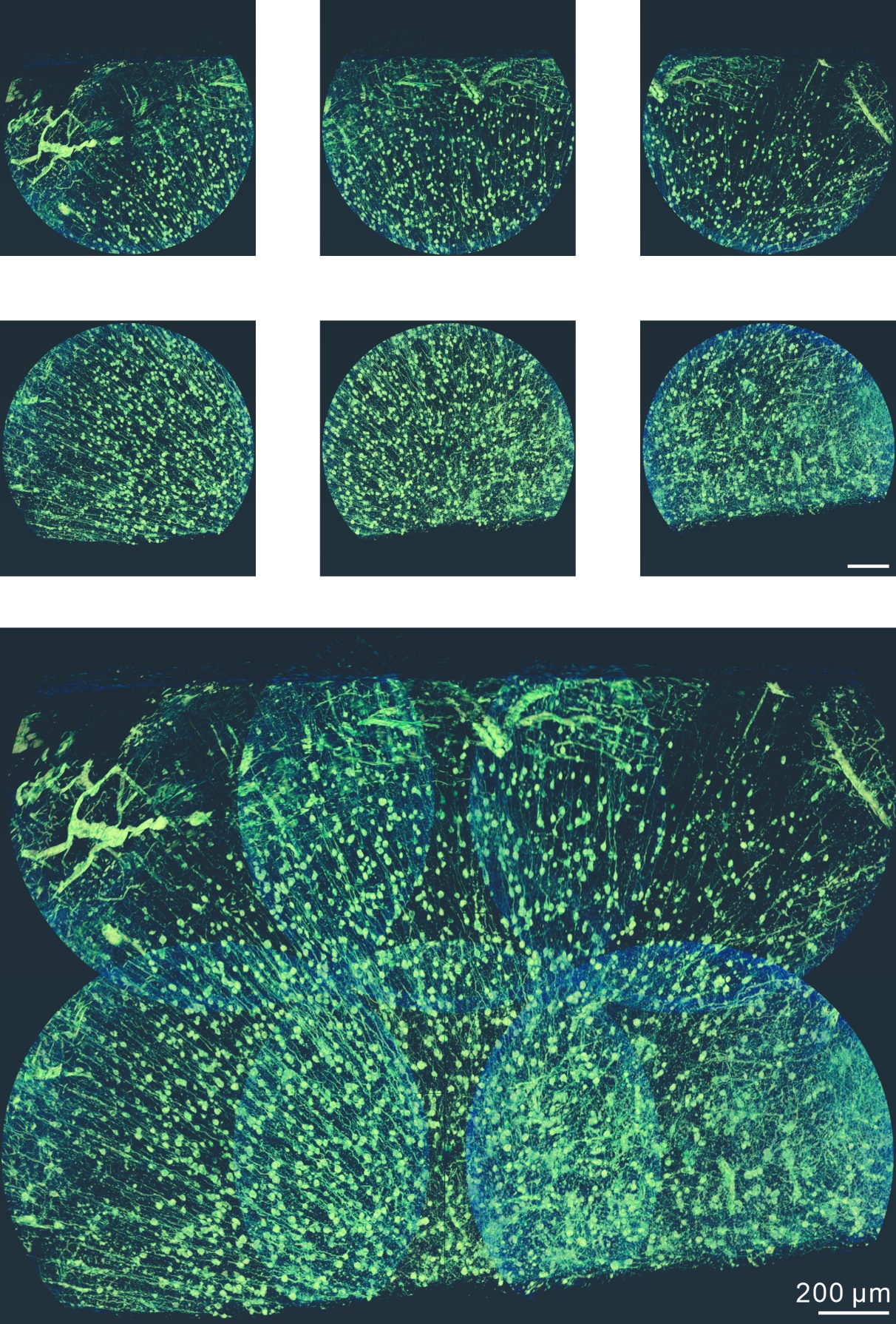

**Supplementary Figure 20. Three-dimensional visualization of the mouse brain cortex (XY plane).** A 3D volume of approximately 9 mm³ was generated by combining six datasets. Scale bar: 200 μm.
